# L-type voltage-gated calcium channel agonists improve hearing loss and modify ribbon synapse morphology in the zebrafish model of Usher Syndrome Type 1

**DOI:** 10.1101/2019.12.16.878231

**Authors:** Alaa Koleilat, Joseph A. Dugdale, Trace A. Christenson, Jeffrey L. Bellah, Aaron M. Lambert, Mark A. Masino, Stephen C. Ekker, Lisa A. Schimmenti

## Abstract

Usher Syndrome (USH) is the most common cause of human deaf/blindness. The zebrafish *myo7aa^-/-^* mutant, faithfully models USH1; homozygous zebrafish are deaf and exhibit circular swimming. We hypothesized that hair cell morphology would differ in *myo7aa^-/-^* mutants compared to wild type. We also tested the hypothesis that agonists of L-type voltage-gated calcium channels would alter ribbon synapse morphology and behavior of zebrafish *myo7aa^-/-^* mutants. We discovered that *myo7aa^-/-^* zebrafish have fewer glutamatergic vesicles tethered to hair cell ribbon synapses, yet maintain a comparable ribbon area. We identified that *myo7aa^-/-^* mutants have fewer total active hair cells, fewer total CTBP2 expressing puncta, and an altered distribution of CTBP2 puncta compared to wildtype. We also identified that *myo7aa^-/-^* mutants have fewer active post-synaptic cells and fewer total MAGUK puncta, compared to wildtype. Behaviorally, *myo7aa^-/-^* mutant fish have abnormal swimming as measured by larger absolute smooth orientations and have little to no acoustic startle. Treatment with L-type voltage-gated calcium channel agonists altered the abnormal cell and behavioral phenotypes toward wildtype. Our data supports that L-type voltage-gated calcium channel agonists induce morphological changes at the ribbon synapse—in both the number of tethered vesicles and the distribution of CTBP2 puncta, shifts swimming behavior towards wildtype swimming and improves acoustic startle response.

**Summary Statement:** We identified that the hair cell biology and behavior of the *myo7aa^-/-^* mutant differs from wildtype and this difference can be rescued using L-type voltage-gated calcium channel agonists.

## INTRODUCTION

A reduction or the complete absence of the ability to detect sounds is the most common impairment of the human senses (Ahmed et al., 2013). About 20 percent of Americans report some degree of hearing loss, making this a major public health concern (Communication) (Prevention, 2016, Prevention, April 1996). Hearing loss is caused by genetic or environmental/infectious factors, with more than fifty percent of hearing loss is attributed to genetics (Morton and Nance, 2006).

Up to 11% of deaf/hard of hearing children and 1 in 6000 people in the United States have Usher Syndrome (USH) (Kimberling et al., 2010). USH is characterized by congenital onset of sensorineural deafness/hard of hearing with later onset of retinitis pigmentosa resulting in significant vision impairment (Usher, 1914). There are three types of USH characterized by degree of hearing, vestibular abnormalities and onset of vision loss (Davenport SLH, 1977).

Usher Syndrome Type I (USH1) is the most severe form of Usher Syndrome with profound bilateral congenital hearing loss and onset of retinitis pigmentosa within the first decade of life. Ten unique loci have been associated with USH1 with variants in *MYO7A* as the most common cause, accounting for 53-63% of affected individuals (Lentz J, 1999). Additionally, pathogenic variants in CDH23, PCDH15, USH1C and USH1G are responsible for 19-35%, 11-19%, 6-7% and 7% respectively (Van Camp G). Each gene encodes structural and motor proteins important for mechanotransduction in the inner ear hair cells (Beurg et al., 2009, Grati and Kachar, 2011, Grillet et al., 2009a, Kazmierczak et al., 2007, Marcotti, 2012, Pepermans and Petit, 2015, Siemens et al., 2004).

Gibson et al. in 1995 identified the first USH locus in the *shaker-1* (*sh1*) mouse. The *shaker-1* mouse presented with hearing loss, head tossing and circling behaviors due to vestibular dysfunction, and upon examination of inner ear hair cells was found to have disorganized stereocilia. Through positional cloning techniques homozygous mutations at the *sh1* locus were identified in *Myo7a*. Two years later, Weil et al. identified that USH1B in humans was also due to pathogenic variants in *MYO7A* (Weil et al., 1997). Ernest et al. in 2000 described a zebrafish model of USH1B caused by a premature stop codon in *myo7aa*, the *mariner* mutant, in which the phenotype of the homozygous recessive larval fish consisted of a circular swimming pattern, defective balance, morphological and functional defects in the inner ear hair cells and most notably the lack of a startle response (Ernest et al., 2000).

*MYO7A* encodes an unconventional actin-binding motor protein, which is important for development and function of the inner ear hair cells. It is specifically involved in upholding the structural integrity of the hair bundle allowing for a mechanical stimulus to be converted into a chemical stimulus. The myosin VIIa protein is localized at the upper tip link density of stereocilia in sensory hair cells (Hasson et al., 1995). Myosin VIIa along with Harmonin and Sans interact with one another to connect the upper end of the tip link (composed of cadherin 23 and protocadherin 15) to the actin cytoskeleton of the stereocilium (Ahmed et al., 2006, Caberlotto et al., 2011, Grati and Kachar, 2011, Grillet et al., 2009b, Siemens et al., 2004). Myosin VIIa is involved in maintaining the tension of the tiplink structure upon positive hair cell deflection. When sound is administered the stereocilia of hair cells are deflected towards the tallest stereocilia allowing for the mechanotransduction channel (MET) located at the apical region of the stereocilia to open (Fig. 1A). The opening of the MET channel causes positively charged cations such as potassium and calcium to flow into the cell and effect depolarization. Once the proper membrane potential is reached, L-type voltage-gated calcium channels (Ca_v_1.3)-localized to the active zones in hair cells adjacent to the ribbon synapse-open, further increasing intracellular calcium concentrations (Brandt et al., 2005, Moser and Vogl, 2016, Sidi et al., 2004). Although calcium has many roles in sensory hair cells, entry of calcium through Ca_v_1.3 is necessary to mediate the release of synaptic vesicles tethered to the ribbon synapse structure (Fuchs, 2005, Moser et al., 2006, Nicolson, 2015, Wong et al., 2014). The ribbon synapse is composed primarily of a central protein, Ribeye encoded by CTBP2, and surrounded by a halo of glutamatergic vesicles (Schmitz et al., 2000). When glutamine is released into the synaptic cleft, it binds onto post-synaptic cell receptors to stimulate cochlear afferents, thus transducing the sound signal (Fig.1A).

**Fig 1.**
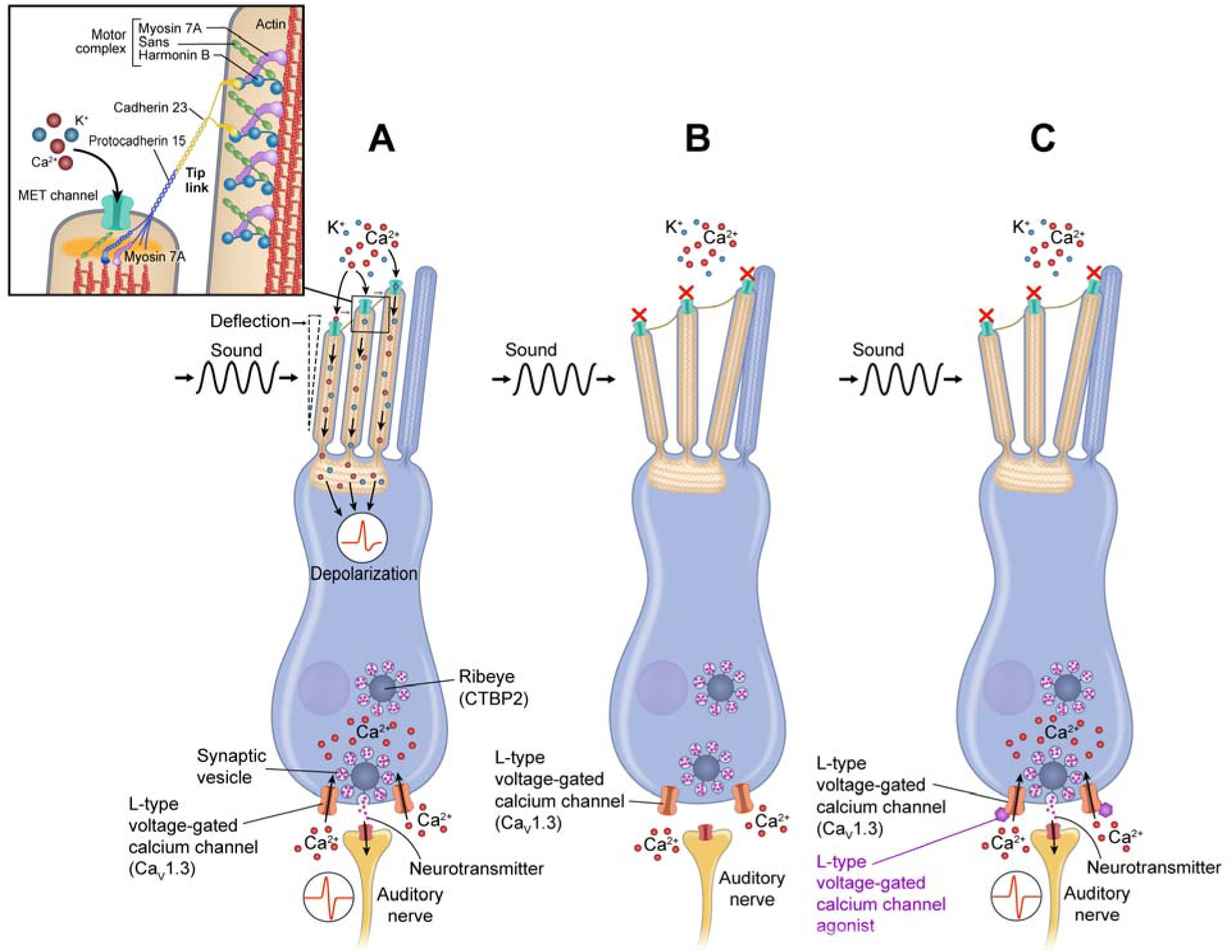
L-type voltage-gated calcium channel agonists may restore function in hair cell. (A) In a normal hair cell, sound causes stereocilia to deflect towards the tallest stereocilia and the mechanotransduction channels (MET) at the top of the stereocilia open in response allowing cations such as calcium and potassium to flow into the cell. This causes a change in membrane potential which leads to L-type voltage gated calcium channels opening at the basolateral sides of the cell. Calcium enters the cell thus increasing intracellular calcium concentrations. Calcium mediates neurotransmitter release from the synaptic vesicles within the ribbon synapse into the synaptic cleft thus stimulating afferent neurons. (B) In cells lacking myo7a expression, proper MET channel gating does not occur and thus the appropriate membrane potential is not reached to allow L-type voltage gated calcium channel to open and thus there is insufficient synaptic transmission to the auditory nerve to create meaningful interactions. We hypothesize that by augmenting the downstream signal in *myo7aa^-/-^* mutant hair cells, we can reconstitute a new functional response to sound by increasing the sensitivity of the calcium channel through treatment with L-type voltage-gated calcium channel agonists (C).

In the absence of myo7a expression, the MET channel is unable to gate properly in response to stereocilia deflections, therefore cell depolarization is not reached and L-type voltage-gated calcium channels do not open, limiting synaptic transmission (Fig. 1B). In this study, we investigated the hypothesis that increasing intracellular calcium in the inner ear hair cell of a *myo7aa^-/-^* mutant would restore hearing and swimming abnormalities as well as change hair cell biology. In order to address this hypothesis, we treated *myo7aa^-/-^* mutants with L-type voltage-gated calcium channel agonists. This class of drugs was previously used in a variety of studies to treat other neurologic conditions, but not hearing loss (Hiramatsu et al., 1997, Kitano et al., 1994, Starkstein et al., 2016).

Here we report that treatment of *myo7aa^-/-^* zebrafish with three separate drugs with L-type voltage-gated calcium channel agonistic activity, specifically (±)-Bay K 8644, Nefiracetam and (R)-Baclofen, can restore hair cell biology and behavior abnormalities observed in untreated mutant zebrafish toward wildtype.

## RESULTS

### Hair cell ribbon synapse ultrastructure, *CTPB2* expression, *MAGUK* expression, and behavior of the myo7aa^-/-^ mutant is different from wildtype

The *myo7aa^-/-^* mutant was first described in 2000 to have absent acoustic startle, balance abnormalities, a deflated swim bladder, and disorganized stereocilia (Ernest et al., 2000); Fig. 2A-D) It was previously reported that the mechanotransduction channel did not uptake the vital dye, FM 1-43, and is therefore inactive in this mutant model (Seiler and Nicolson, 1999); Fig. 2A, B). To elucidate the ultrastructural state of the ribbon synapse structure in hair cells we used transmission electron microscopy (TEM) (Fig. 2E, F). The mean ± standard error of the mean of wildtype ribbon area is 37866.7 ± 3166.2 nm^2^ (n=35 ribbons; 6 zebrafish) and the mean of the *myo7aa^-/-^* mutants is 30992.6 ± 3347.2 nm^2^ (n=35 ribbons; 5 zebrafish); therefore the ribbon area between *myo7aa^-/-^* mutants and wildtype larvae is comparable (Fig. 2G, t-test, p=0.14). The mean number of tethered vesicles for wildtype is 19 ± 1 vesicles (n=32 ribbons, 6 zebrafish) which is statistically different from the mean for the *myo7aa^-/-^* mutants 15 ± 1 vesicles (n=34 ribbons, 5 zebrafish) (Fig. 2H, t-test, p<0.01).

**Fig. 2.**
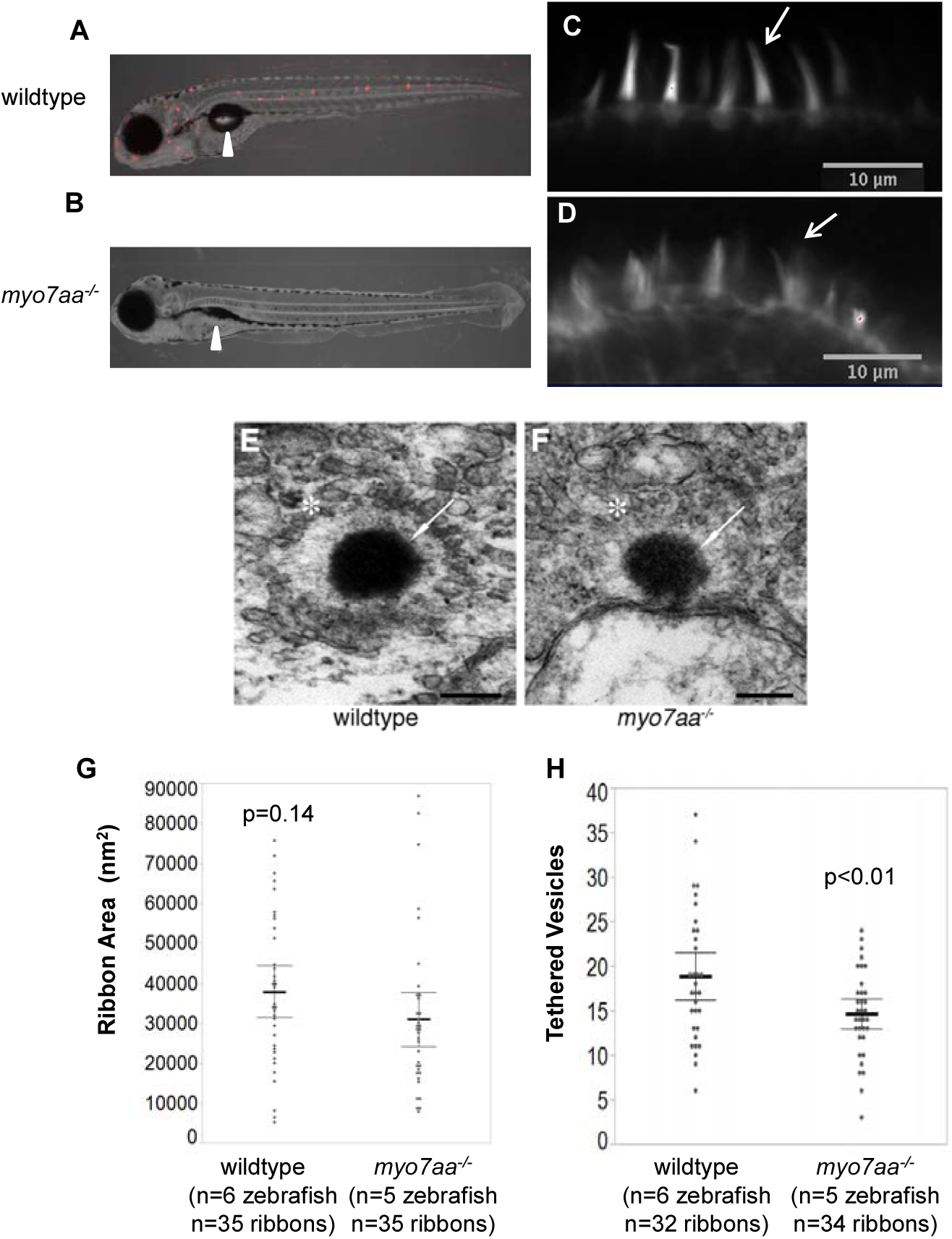
*myo7aa^-/-^* larvae at 5 dpf have altered mechanotransduction activity, stereocilia structure and ribbon synapse structure. (A, B) Representative lateral view of 5 dpf wildtype larvae (A) and *myo7aa^-/-^* larvae (B) briefly exposed to FM1-43 obtained using Light Sheet Fluorescence microscopy. Labeling with FM1-43 indicates functional mechanotransduction. White arrowhead points to the swimbladder; *myo7aa^-/-^* larvae do not have an inflated swimbladder organ. (C, D) Representative images of 5 dpf wildtype stereocilia (C) and *myo7aa^-/-^* (D) stained with Alexa Fluor 488 Phalloidin, a high-affinity filamentous actin probe, obtained using confocal microscopy. White arrow highlights organized and smooth stereocilia in wildtype hair cells and disorganized and splayed stereocilia in myo7aa^-/-^ hair cells. Note: not all stereocilia are splayed in *myo7aa^-/-^* hair cells. (E, F) Representative images of 5dpf wildtype ribbon synapse structure (E) and myo7aa^-/-^ ribbon synapse (F) obtained with transmission electron microscopy. White arrow points to ribeye protein and white star indicates a halo of tethered vesicles. Scale bars: 200 nm. (G) Wildtype ribbon synapses have a comparable ribbon area to *myo7aa^-/-^* ribbon synapses (t-test). (H) Wildtype ribbon synapses have an increased number of tethered vesicles compared to *myo7aa^-/-^* ribbon synapses (t-test). Black bold line represents the mean of the data set and error bars are 95% confidence intervals.

We further explored the ribbon synapse in the *myo7aa^-/-^* mutants and performed immunohistochemistry to assess differences in pre- and post-synaptic markers of the hair cells. The ribeye protein is the result of alternative splicing of the transcription factor CTBP2. Ribeye has two main roles: clustering of the Ca_v_1.3 channels at the presynapse and stabilizing contacts with the afferent neurons (Fig. 3A) (Lv et al., 2016, Sheets et al., 2011). CTBP2 puncta can be observed in hair cells and we define an active hair cell as any hair cell with at least one CTBP2 puncta. We assessed CTBP2 expression in zebrafish lateral line hair cells and quantified the number of active hair cells, total puncta and the distribution of puncta. The wildtype neuromasts had a mean of 19 ± 1 active hair cells (n=22 neuromasts, n=18 zebrafish) and a mean of 62 ± 4 total CTBP2 puncta per neuromast (n=22 neuromasts, n=17 zebrafish). These two parameters are statistically significant compared to the *myo7aa^-/-^* mutants with a mean of 12 ± 1 active hair cells (n=18 neuromasts, n=17 zebrafish) and a mean of 34 ± 3 total CTBP2 puncta per neuromast (n=18 neuromasts, n=17 zebrafish) (t-test, p<0.0001, p<0.0001) indicating that there are fewer active hair cells and fewer CTBP2 puncta per neuromast in the *myo7aa^-/-^* mutants (Fig. 3D, E). We also investigated the distribution of CTBP2 puncta and identified that most hair cells from the *myo7aa^-/-^* mutant neuromasts had two puncta per cell whereas most wildtype hair cells had three puncta per cell (Fig. 3F). Additionally, the proportion of hair cells with two CTBP2 puncta and three CTBP2 puncta are statistically significant between wildtype and *myo7aa^-/-^* mutants accompanied by no differences in the proportion of 1, 4, 5 and 6+ CTBP2 puncta between the two groups (Table S2).

**Fig. 3.**
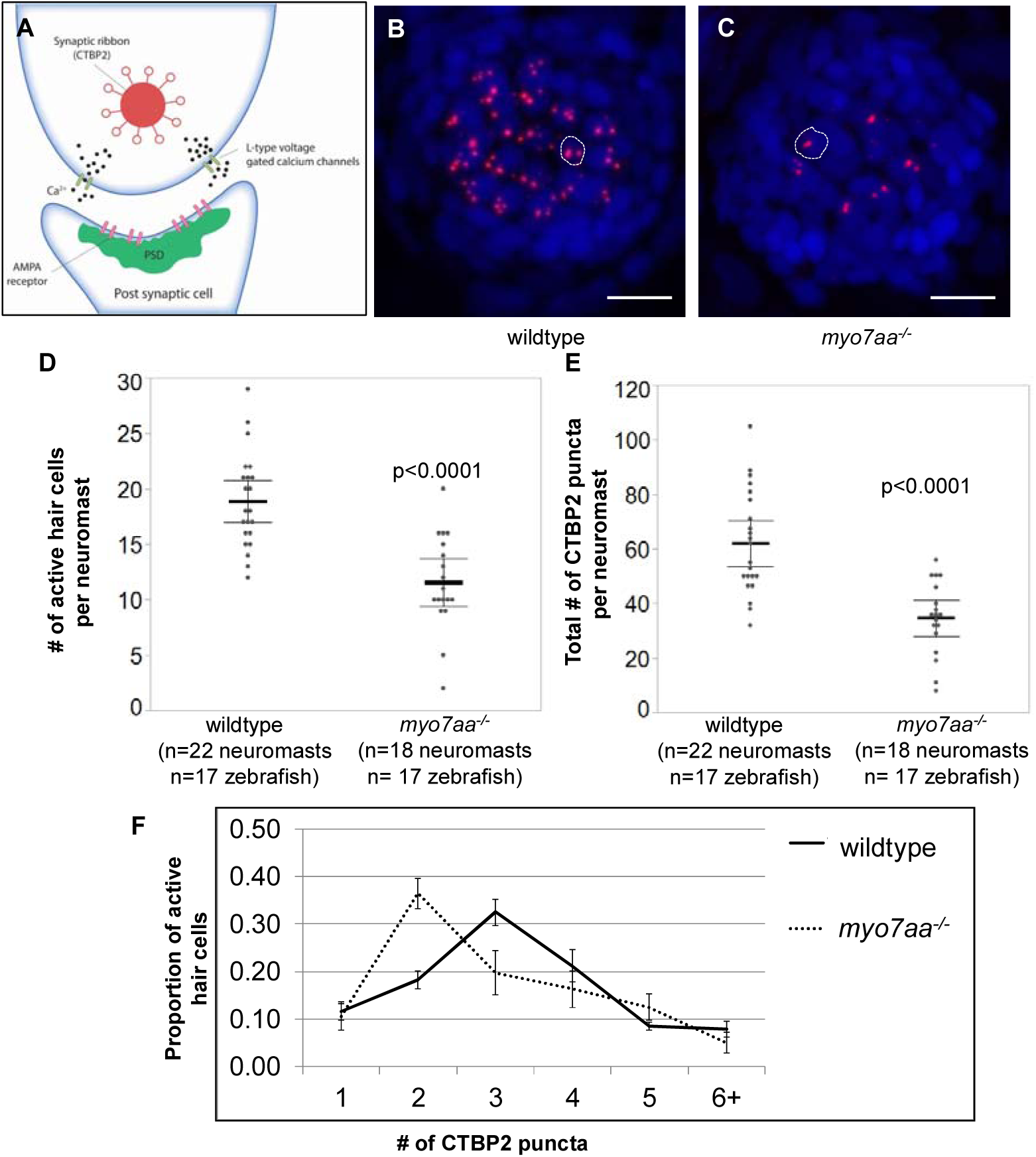
*myo7aa^-/-^* CTBP2 expression in 5 dpf larvae is different from 5 dpf wildtype larvae. (A) Cartoon depiction of the basal end of a hair cell innervated by the auditory nerve. CTBP2 is alternatively spliced to produce ribeye protein, which is a component of the synaptic ribbon. Ribeye has a halo of tethered vesicles which forms the synaptic ribbon. (B, C) Representative maximum intensity projection (z-stack top-down image) of neuromast from 5dpf wildtype larvae (B) and *myo7aa^-/-^* larvae (C) labeled with DAPI labeled nuclei (blue) and Ribeye (red). The dotted circle outlines one hair cell. Scale bars: 10 µm. (D) 5dpf wildtype neuromasts have a greater number of active hair cells compared to *myo7aa^-/-^* neuromasts. An active hair cell is defined as a hair cell with at least one CTBP2 puncta (t-test). (E) 5dpf wildtype neuromasts have a greater number of total CTBP2 puncta to *myo7aa^-/-^* neuromasts (t-test). Black bold line represents the mean of the data set and error bars are 95% confidence intervals. (F) Further analysis of the distribution of CTBP2 puncta across a collection of active hair cells reveals that most 5dpf wildtype active hair cells contain 2 CTBP2 puncta compared to 3 CTBP2 puncta in *myo7aa^-/-^* hair cells. All error bars are 95% confidence intervals.

We also explored any differences in the post-synaptic hair cell marker, MAGUK. Membrane associated guanylate kinase, MAGUK, was used to assess afferent post-synaptic structures in the afferent fiber terminal (Anderson, 1996, Lv et al., 2016, Sheets et al., 2011). This kinase family targets and anchors glutamate receptors to the synaptic terminal on the post-synaptic cell as part of the postsynaptic density (PSD) (Fig. 4A). Wildtype neuromasts had a mean of 22 ± 1 active post-synaptic cells (n=29 neuromasts, n=29 zebrafish) and a mean of 59 ± 3 total MAGUK puncta per neuromast (n=29 neuromasts, n=29 zebrafish). These two parameters are statistically significant compared to the *myo7aa^-/-^* mutants which have a mean of 16 ± 1 active post-synaptic cells (n=23 neuromasts, n=23 zebrafish) and a mean of 46 ± 2 total MAGUK puncta per neuromast (n=23 neuromasts, n=23 zebrafish) (t-test, p<0.0001, p=0.001). This indicates that there are fewer active post-synaptic cells and fewer MAGUK puncta per neuromast in the *myo7aa^-/-^* mutants (Fig. 4D, E). We also investigated the distribution of MAGUK puncta and identified that most post-synaptic cells from the *myo7aa^-/-^* mutant neuromasts had 3 puncta per cell whereas most wildtype hair cells had either 2 or 3 puncta per cell; however, this difference is not statistically significant (Fig. 4F, Table S2)

**Fig. 4.**
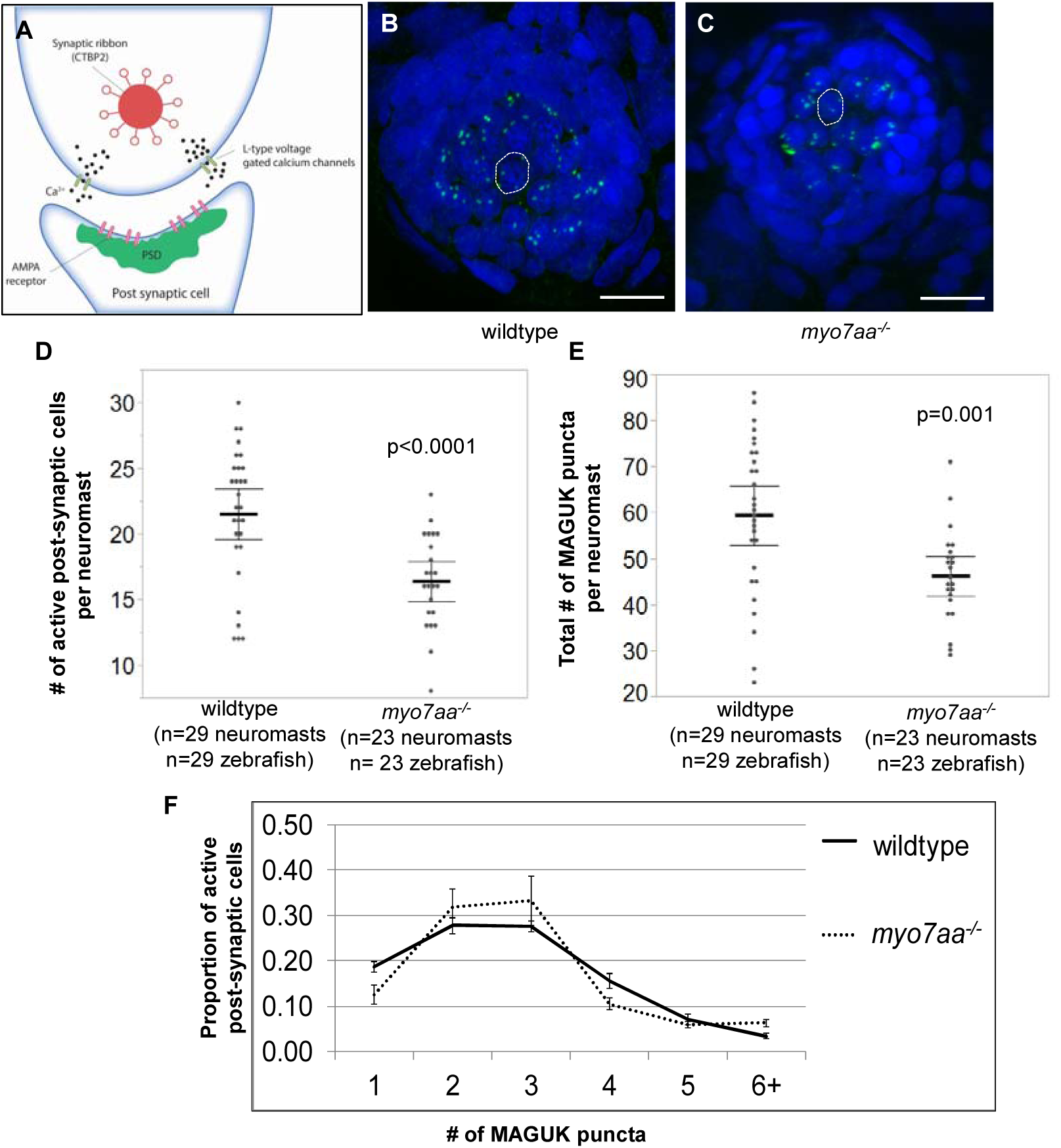
*myo7aa^-/-^* MAGUK expression in 5 dpf larvae is different from 5 dpf wildtype larvae. (A) Cartoon depiction of the basal end of a hair cell innervated by the auditory nerve. MAGUK (membrane associated guanylate kinases) are a kinase family that target and anchor glutamate receptors to the synaptic terminals on the post-synaptic cell as part of the postsynaptic density (PSD). (B, C) Representative maximum intensity projection (z-stack top-down image) of neuromast from 5 dpf wildtype larvae (B) and *myo7aa^-/-^* larvae (C) labeled with DAPI labeled nuclei (blue) and MAGUK (green). The dotted circle outlines one hair cell. Scale bars: 10 µm. (D) 5dpf wildtype neuromasts have a greater number of active post-synaptic cells compared to *myo7aa^-/-^* neuromasts. An active post-synaptic cell is defined as a cell with at least one MAGUK puncta (t-test). (E) 5dpf wildtype neuromasts have a greater number of total MAGUK puncta to *myo7aa^-/-^* neuromasts (t-test). Black bold line represents the mean of the data set and error bars are 95% confidence intervals. (F) Further analysis of the distribution of MAGUK puncta across a collection of active hair cells reveals that most 5 dpf wildtype active hair cells contain either 2 or 3 MAGUK puncta compared to 3 MAGUK puncta in *myo7aa^-/-^* hair cells. All error bars are 95% confidence intervals.

The abnormal circular and loop-like swimming behavior of the *myo7aa^-/-^* mutant is apparent by observation using a tactile stimulus to the tail or through spontaneous swimming. We were able to quantify this abnormal swimming behavior by using a previously described behavior assessment (Branson et al. 2009). We quantified the absolute smooth orientation or global change in body orientation as a function of time. The absolute smooth orientation of wildtype fish was 295 ± 5° (n=46 zebrafish) compared to the *myo7aa^-/-^* mutant larvae which had a mean absolute smooth orientation of 486 ± 5° (n=61 zebrafish)(t-test; p<0.0001) (Fig. 5).

**Fig. 5.**
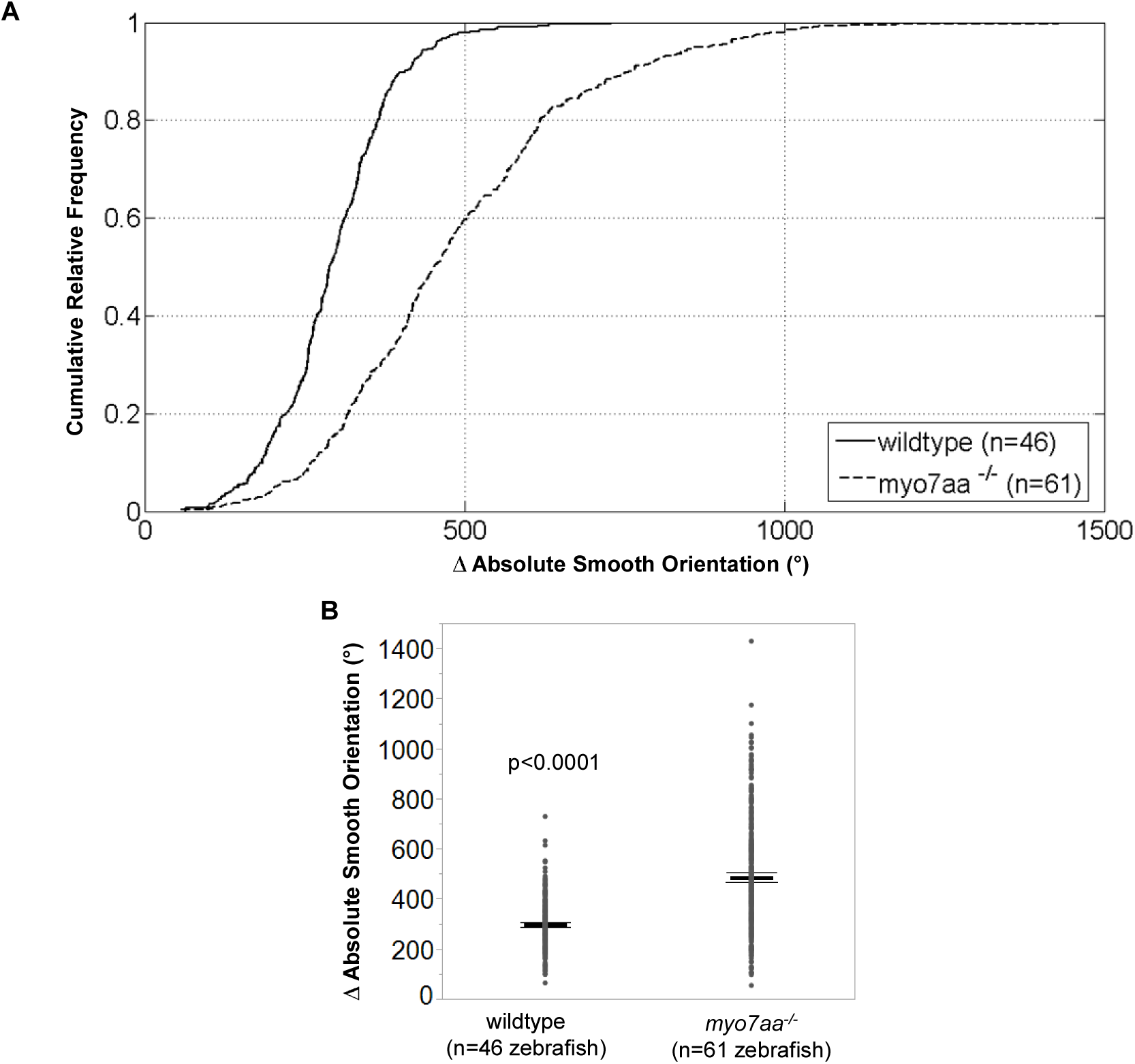
5 dpf *myo7aa^-/-^* larvae exhibit larger turning angles. Movement tracking of 5dpf wildtype and *myo7aa^-/-^* larvae over a 2.5 minute interval with a 5 ms electric stimulus (50 mV) administered every 20 s. Ctrax software was used for video processing and Matlab 2012b for video analysis. (A, B) *myo7aa^-/-^* larvae have larger absolute smooth orientations (turning angles) compared to wildtype larvae. Individual turning angles from a population of wildtype and *myo7aa^-/-^* larvae were used to construct the lines (t-test). Black bold line represents the mean of the data set and error bars are 95% confidence intervals.

### L-type voltage gated calcium channel agonists alter myo7aa^-/-^ ribbon synapse ultrastructure

Through transmission electron microscopy we established that ribbon synapse structures differ between 5 days post fertilization (dpf) wildtype and *myo7aa^-/-^* mutant fish, specifically the number of vesicles tethered to the electron-dense structure, ribeye. To explore whether L-type voltage-gated calcium channel agonists can modify hair cell ribbon synapse components we incubated 4 dpf *myo7aa^-/-^* mutant embryos overnight in either 5 μM (±)-Bay K 8644, 250 μM Nefiracetam or 125 μM (R)-Baclofen and performed transmission electron microscopy on 5 dpf. We observed that the mean area of the electron dense structure, ribeye, for *myo7aa^-/-^* mutant fish incubated with either 5 μM (±)-Bay K 8644 or 250 μM Nefiracetam was 27274.97 ± 2231.61 nm^2^ (n=35 ribbons, n=5 zebrafish)(t-test, p=0.36) and 24024.14 ± 2335.09 nm^2^ (n=29 ribbons, n=4 zebrafish)(t-test, p=0.09), respectively, and thus did not alter the area of the ribbon synapse. Additionally, neither drug altered the number of vesicles tethered to ribeye with a mean of 17 ± 1 (n=35 ribbons, n=5 zebrafish) (t-test, p=0.07) and a mean of 14 ± 1 (n=29 ribbons, n=4 zebrafish) (t-test, p=0.54), respectively. However, treatment with 125 μM (R)-Baclofen did modify both parameters with a mean area of 44071.6 ± 3008 nm^2^ (n=50 ribbons, n=5 zebrafish) (t-test, p=0.02) and 17 ± 1 tethered vesicles (n=46 ribbons, n=5 zebrafish) (t-test, p=0.03) (Fig. 6). None of the three drugs showed any adverse effect on wildtype ribbon synapse ultrastructure compared to untreated wildtype (Fig. S1).

**Fig. 6.**
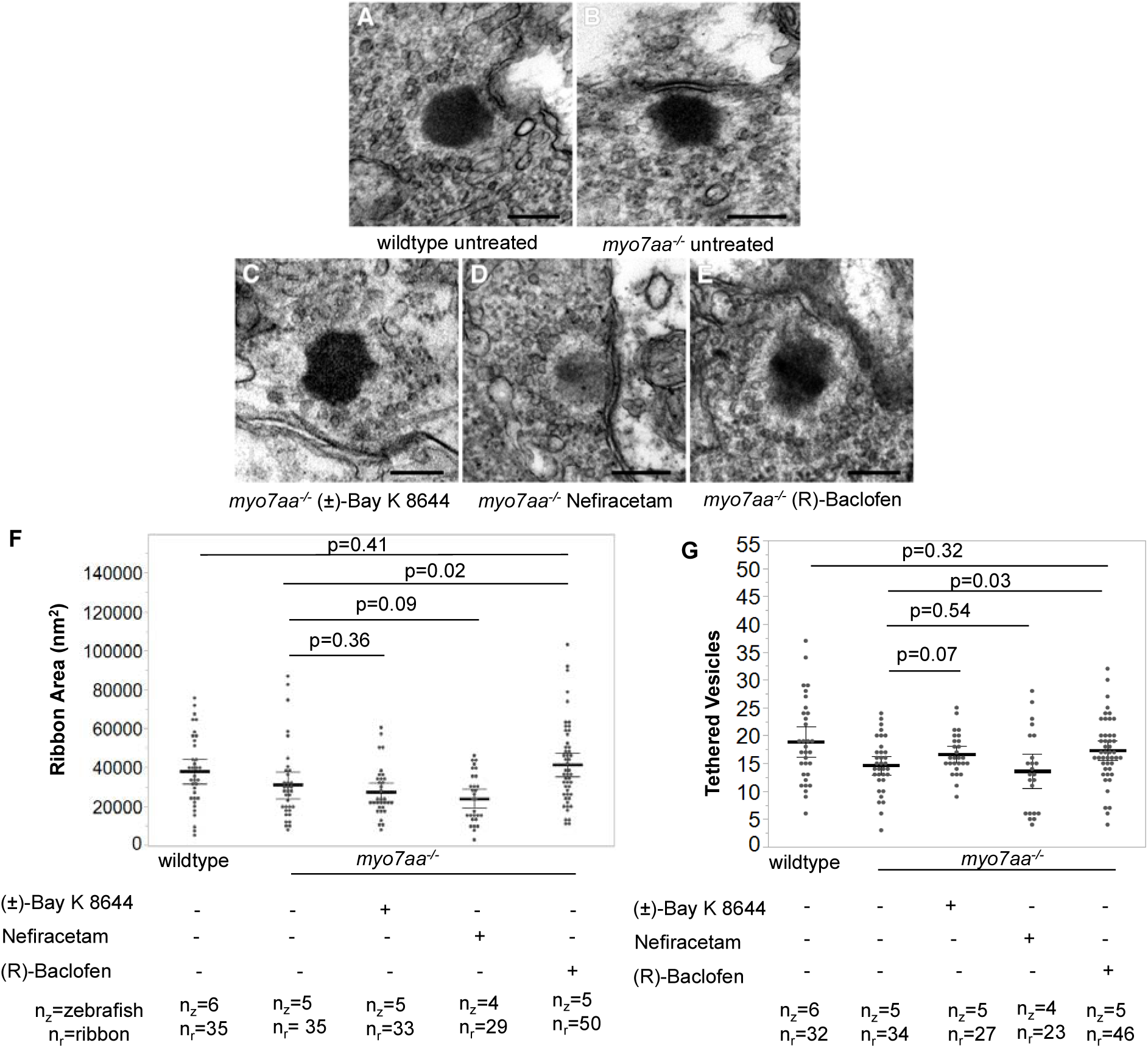
*myo7aa^-/-^* mutant ribbon synapse abnormalities diminished with exposure to (R)-Baclofen. (A-E) Representative images of 5dpf wildtype untreated ribbon synapse structure (A) myo7aa^-/-^ untreated ribbon synapse (B) and *myo7aa^-/-^* treated with 5 µM (±)-Bay K 8644 (C), 250 µM Nefiracetam (D) and 125 µM (R)-Baclofen (E) obtained with transmission electron microscopy. Scale bars: 1000 nm. (F) *myo7aa^-/-^* untreated ribbon synapse area is smaller compared to myo7aa^-/-^ treated with 125 µM (R)-Baclofen. Additionally, wildtype untreated ribbon synapses have a comparable ribbon area to myo7aa^-/-^ treated with 125 µM (R)-Baclofen ribbon synapses (t-test). (G) *myo7aa^-/-^* untreated ribbons have fewer tethered vesicles compared to myo7aa^-/-^ treated with 125 µM (R)-Baclofen. Wildtype untreated ribbons have a comparable number of tethered vesicles to *myo7aa^-/-^* treated with 125 µM (R)-Baclofen ribbon synapses (t-test). This indicates that there is a rescue in the number of tethered vesicles upon treatment with (R)-Baclofen. Black bold line represents the mean of the data set and error bars are 95% confidence intervals.

### L-type voltage-gated calcium channel agonists alter CTBP2 and MAGUK expression

We previously identified that the *myo7aa^-/-^* mutant has different CTBP2 and MAGUK gene expression compared to wildtype. We discovered that *myo7aa^-/-^* mutants have fewer active hair cells and post-synaptic cells, fewer total CTBP2 and MAGUK puncta and a different distribution of CTBP2 puncta compared to wildtype. Here we explored the effect of L-type voltage-gated calcium channel agonists on CTBP2 and MAGUK expression. We identified that wildtype larvae treated with either 250 μM Nefiracetam or 125 μM (R)-Baclofen did not alter the overall distribution of CTBP2 puncta, however incubation with 5 μM (±)-Bay K 8644 did shift the curve so that most hair cells had 2 puncta per cell compared to 3 puncta per cell in the untreated groups (Fig. 7 A-D, I; Table S3). There was generally no change in the distribution of MAGUK puncta with incubation of either 5 μM (±)-Bay K 8644, 250 μM Nefiracetam or 125 μM (R)-Baclofen (Fig. 7E-H, J; Table S4). Additionally, there was an increase in the total number of active hair cells per neuromast and the total number of CTBP2 puncta per neuromast for wildtype fish incubated in 5 μM (±)-Bay K 8644, but no change in both parameters upon incubation with either 250 μM Nefiracetam or 125 μM (R)-Baclofen (Fig. S2A). There was, however, an increase in total number of active post-synaptic cells per neuromast and the total number of MAGUK puncta per neuromast for wildtype fish incubated in 5 μM (±)-Bay K 8644 or 125 μM (R)-Baclofen, but no change with 250 μM Nefiracetam (Fig. S3A). We also identified that *myo7aa^-/-^* mutant larvae treated with either 5 μM (±)-Bay K 8644, 250 μM Nefiracetam or 125 μM (R)-Baclofen did alter the distribution of CTBP2 puncta so that most hair cells had 3 puncta per cell (Fig. 7K-N, S; Table S3). There was also generally no statistically significant change in the distribution of MAGUK puncta with incubation of either 5 μM (±)-Bay K 8644, 250 μM Nefiracetam or 125 μM (R)-Baclofen, (Fig. 7O-R, T; Table S4). Moreover, there was an increase in the total number of active hair cells per neuromast, total number of CTBP2 puncta per neuromast, total number of post-synaptic cells per neuromast and total number of MAGUK puncta per neuromast for *myo7aa^-/-^* fish incubated in 5 μM (±)-Bay K 8644, but no change in any of the parameters upon incubation with either 250 μM Nefiracetam or 125 μM (R)-Baclofen (Fig. S2B, Fig. S3B).

**Fig 7.**
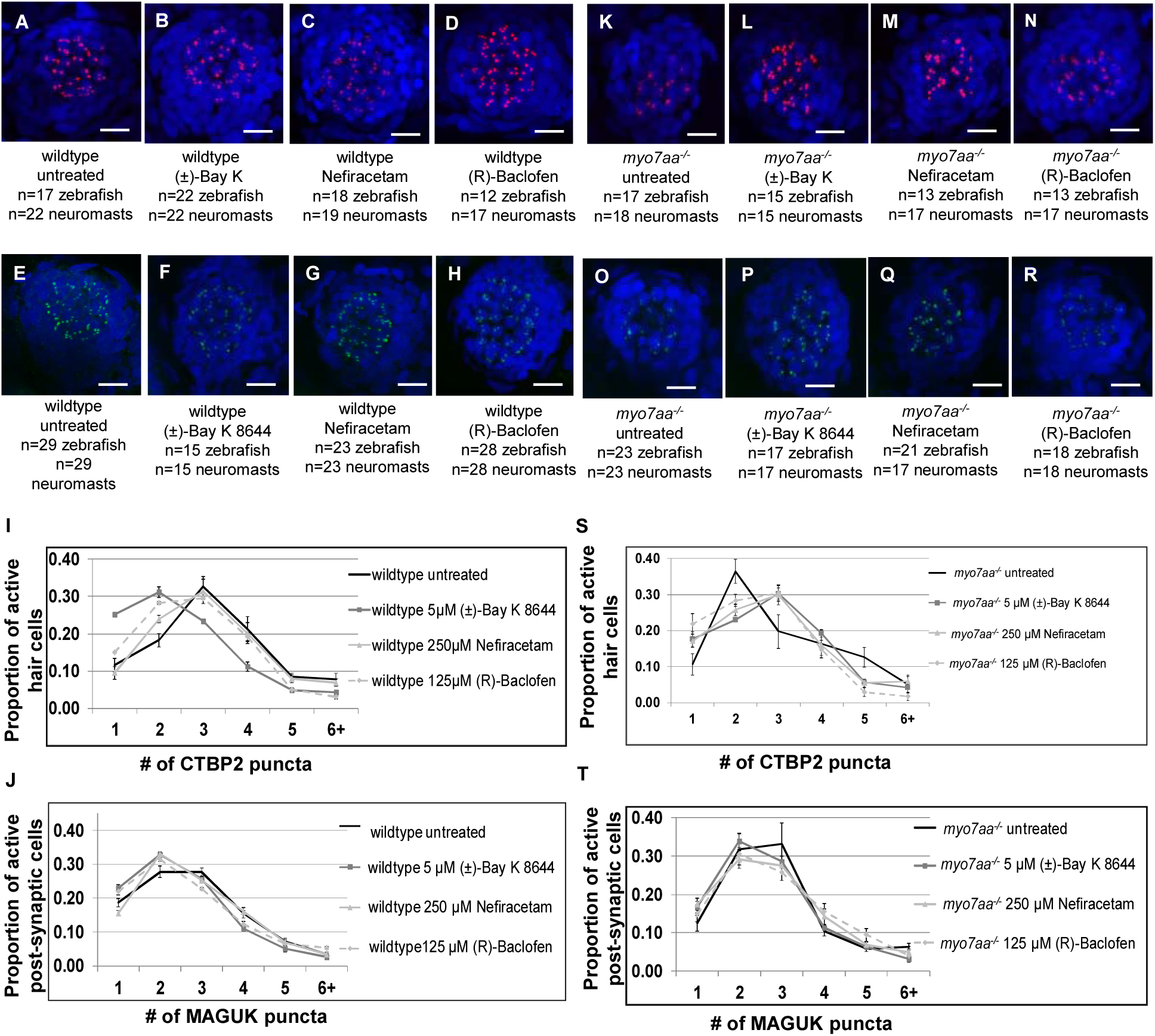
Distribution of CTBP2 and MAGUK puncta more closely resembles wildtype distribution with exposure to L-type voltage-gated calcium channel agonists in 5dpf *myo7aa^-/-^* neuromasts. (A-D, J-N) Representative maximum intensity projection (z-stack top-down image) of neuromast from 5 dpf wildtype larvae untreated and treated (A-D) and *myo7aa^-/-^* larvae untreated and treated (K-N) labeled with DAPI labeled nuclei (blue) and Ribeye (red). Scale bars: 10 µm. (I) A majority of 5 dpf untreated wildtype active hair cells have 3 CTBP2 puncta. Upon incubation with 5 µM (±)-Bay K 8644 a majority of active hair cells shifted to 2 CTBP2 puncta. There was no change with incubation of either 250 µM Nefiracetam or 125 µM (R)-Baclofen. (S) A majority of 5dpf untreated *myo7aa^-/-^* active hair cells have 2 CTBP2 puncta. Upon incubation with 5 µM (±)-Bay K 8644, 250 µM Nefiracetam or 125 µM (R)-Baclofen a majority of hair cells have 3 CTBP2 puncta. (E-H, O-R) Representative maximum intensity projection (z-stack top-down image) of neuromast from 5 dpf wildtype larvae untreated and treated (E-H) and *myo7aa^-/-^* larvae untreated and treated (O-R) labeled with DAPI labeled nuclei (blue) and Ribeye a (red). Scale bars: 10 µm. (J) A majority of 5 dpf untreated wildtype active hair cells have 2 or 3 MAGUK puncta. Upon incubation with 5 µM (±)-Bay K 8644, 250 µM Nefiracetam or 125 µM (R)-Baclofen a majority of active hair cells have 2 MAGUK puncta. (T) A majority of 5 dpf untreated *myo7aa^-/-^* active hair cells have 3 MAGUK puncta. Upon incubation with 5 µM (±)-Bay K 8644, 250 µM Nefiracetam or 125 µM (R)-Baclofen a majority of hair cells shift to 2 MAGUK puncta. All error bars are 95% confidence intervals.

### L-type voltage-gated calcium channel agonists restore myo7aa^-/-^ abnormal swimming

To determine the effect of L-type voltage-gated calcium channel agonists on swimming behavior in the *myo7aa^-/-^* mutants, we used a behavior assessment that quantifies turning angles as a function of time (absolute smooth orientation). Wildtype fish incubated in 5 μM (±)-Bay K 8644 had smaller absolute smooth orientations with a mean of 218 ± 6° (n=15 zebrafish) compared to untreated wildtype larvae with a mean of 295 ± 5° (n=46 zebrafish), however swimming traces were similar (t-test; p<0.0001) (Fig. 8A, B, C). Incubation with 250 μM Nefiracetam or 125 μM (R)-Baclofen had similar absolute smooth orientations compared to untreated wildtype fish 309 ± 10° (n=17 zebrafish) and 313 ± 13° (n=11) and swimming trajectories, respectively (t-test, p=0.22, p=0.23) (Fig. 8A, B, C). *Myo7aa^-/-^* mutant fish incubated in 5 μM (±)-Bay K 8644, 250 μM Nefiracetam or 125 μM (R)-Baclofen showed improved swimming behavior with decreased absolute smooth orientation 252 ± 16° (n=11), 450 ± 13° (n=22), 451 ± 11° (n=20) compared to untreated controls, respectively (t-test, p<0.0001, p=0.03, p=0.01) (Fig.8A, C). *Myo7aa^-/-^* fish incubated with any of the three drugs displayed smoother swimming trajectories with less circular swimming and loop-like swimming and more zigzag swimming (Fig. 8B). Dose response and time response experiments were performed for Nefiracetam and (R)-Baclofen on wildtype and *myo7aa^-/-^* mutants (Fig. S4-7). Three doses of Nefiracetam were tested, 85 μM, 125 μM and 250 μM. Wildtype absolute smooth orientations did not differ from untreated fish except at 125 μM and *myo7aa^-/-^* mutant absolute smooth orientations did not differ from untreated fish except at 250 μM (Fig. S4). We also tested if 1 hour incubation would suffice to see any effect and we concluded that there was no difference in either group after 1 hour incubation (Fig. S5). Five doses of (R)-Baclofen were tested, 75 μM, 125 μM, 250 μM, 500 μM and 1 mM. Wildtype absolute smooth orientations did not differ from untreated fish except at 75 μM, 250 μM, 500 μM and *myo7aa^-/-^* mutant absolute smooth orientations did differ from untreated fish at all doses (Fig. S6). We also tested a shorter incubation time at the 125uM dose and identified that a 1 hour incubation of (R)-Baclofen is sufficient to see an effect in the *myo7aa^-/-^* mutants (Fig. S7). There was no difference between wildtype untreated and treated with 125 μM (R)-Baclofen after the 1 hour incubation (Fig. S7).

**Fig. 8.**
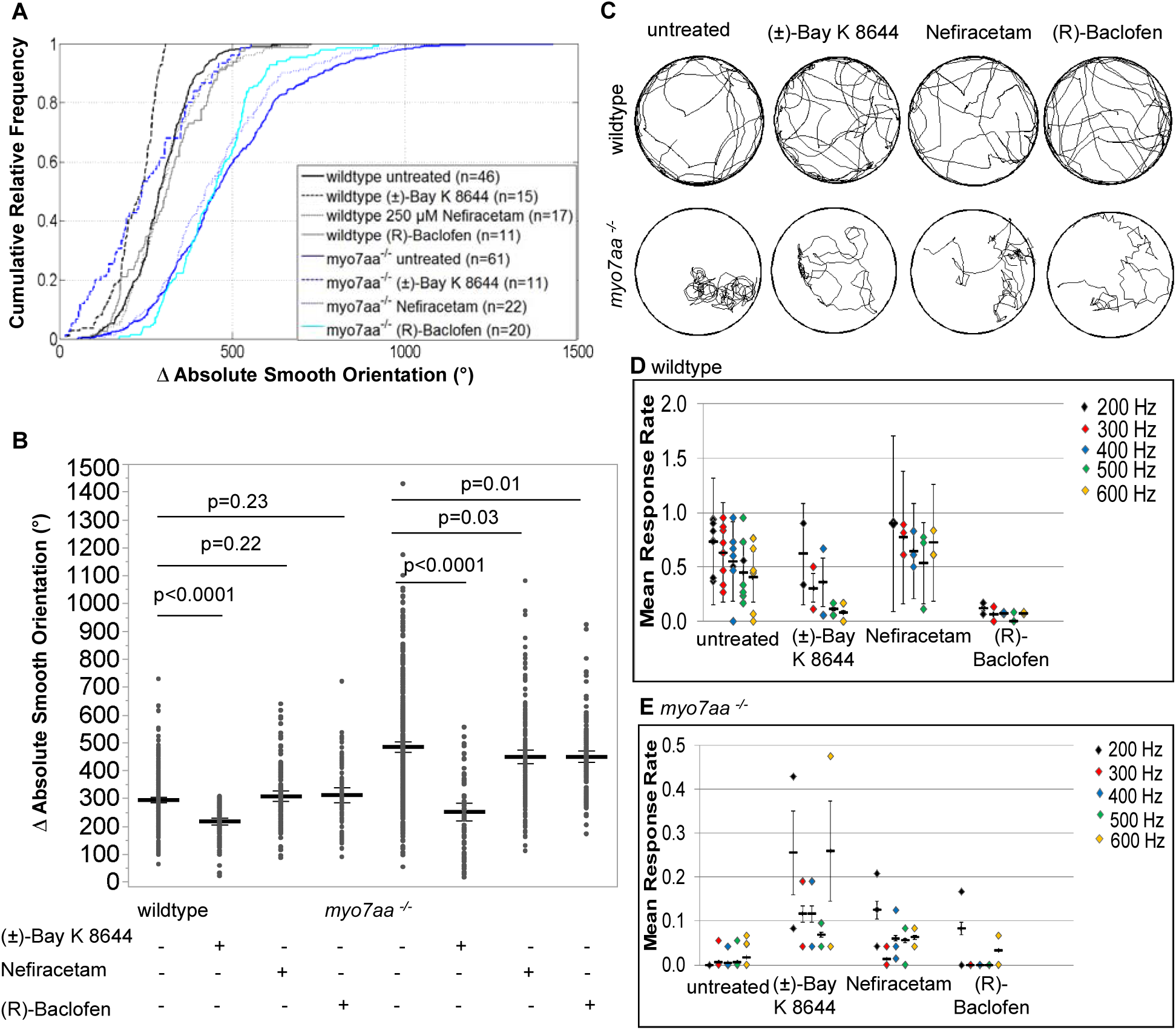
*myo7aa^-/-^* swimming behavior and acoustic startle response improved with L-type voltage-gated calcium channel agonists. Movement tracking of 5 dpf wildtype and *myo7aa^-/-^* larvae over a 2.5 minute interval with a 5 ms electric stimulus (50 mV) administered every 20 s. Ctrax software was used for video processing and Matlab 2012b for video analysis. (A, B) 250 µM Nefiracetam and 125 µM (R)-Baclofen did not affect the absolute smooth orientation of wildtype larvae, however, 5 µM (±)-Bay K 8644 resulted in a decreased absolute smooth orientation compared to untreated wildtype larvae. *Myo7aa^-/-^* larvae treated with 5 µM (±)-Bay K 8644, 250 µM Nefiracetam and 125 µM (R)-Baclofen individually all gave decreased absolute smooth orientations, with the most robust response observed in 5 µM (±)-Bay K 8644 incubation. Individual turning angles from a population of wildtype and *myo7aa^-/-^* larvae were used to construct the lines (t-test). Black bold line represents the mean of the data set and error bars are 95% confidence intervals. (C) Sample movement traces of wildtype and *myo7aa^-/-^* larvae in individual treatment groups indicates that in the presence of 5 µM (±)-Bay K 8644, 250 µM Nefiracetam and 125 µM (R)-Baclofen *myo7aa^-/-^* larvae swimming changes to include smoother trajectories with more zig-zag like swimming and fewer circling episodes and overall movement traces of wildtype in the 4 treatments is unaffected. Diameter of the wells is 20 mm. (D, E) Acoustic startle response was captured by administering 3 stimuli at each frequency per experiment. Videos were scored blindly and the mean number of responses per total stimuli (mean response rate) was determined. 5dpf untreated wildtype larvae respond most at 200 Hz and *myo7aa^-/-^* larvae have little to no response at all frequencies. Upon incubation with 5 µM (±)-Bay K 8644 wildtype acoustic startle decreased at all frequencies, however *myo7aa^-/-^* larvae had a significant increase with over 20% response rate at 200 and 600 Hz. Incubation in 250 µM Nefiracetam increased acoustic startle in wildtype and *myo7aa^-/-^* larvae at all frequencies, although the increase in the *myo7aa^-/-^* larvae is modest. 125 µM (R)-Baclofen decreased acoustic startle response significantly in wildtype larvae and had little to no effect on acoustic startle in the *myo7aa^-/-^* larvae. Black bold line represents the mean of the data set and error bars are mean variance.

Lastly, we assessed if there was a change in swim bladder inflation upon incubation in the *myo7aa^-/-^* mutants with 250 μM Nefiracetam and 125 μM (R)-Baclofen and did not see any change (Fig. S8).

### L-type voltage-gated calcium channel agonists increase myo7aa^-/-^ acoustic startle response

The *myo7aa^-/-^* mutant is profoundly deaf and has little to no acoustic startle to sounds at 200, 300, 400, 500 or 600 Hz (Fig.8E), however a profound deafness does not equate to an abolished ability to detect sounds if in the right environment given the correct cellular machinery. We assessed whether L-type voltage-gated calcium channel agonists can improve the ability of *myo7aa^-/-^* mutants to startle to an acoustic stimulus. Wildtype fish untreated responded most frequently at 200 Hz with a mean response rate of 0.70 and the lowest mean response rate at 600 Hz. Wildtype fish treated with 5 μM (±)-Bay K 8644 had decreased responses at all frequencies, except 200 Hz (Table S5). Incubation with 125 μM (R)-Baclofen also decreased acoustic startle responses at all frequencies, however this may be due to the sedation effect of this drug (Finnimore et al., 1995, Renier et al., 2007) (Fig. 8D, Table S5). Interestingly, both 5 μM (±)-Bay K 8644 and 250 μM Nefiracetam increased acoustic startle response in the *myo7aa ^/-^* mutant fish, however, (±)-Bay K 8644 had a more robust effect and increased responses at all frequencies. We did not see any improvement of *myo7aa^-/-^* mutant acoustic startle response upon incubation with 125 μM (R)-Baclofen (Fig. 8E, Table S5).

## DISCUSSION

The zebrafish model of USH1 was first characterized in 2000 and shows phenotypes similar to human patients (Ernest et al., 2000). The *myo7aa^-/-^* mutants are deaf at the onset of hearing (5 dpf), have vestibular abnormalities manifested in abnormal circular and loop like swimming, have irregular stereocilia and do not uptake the FM1-43 vital dye, indicating an inactive MET channel. In the present study we identified that 1) the *myo7aa^-/-^* mutants have comparable areas of the ribeye protein in the ribbon synapse and significantly fewer tethered vesicles; 2) the *myo7aa^-/-^* mutants have significantly fewer numbers of active hair cells, fewer total CTBP2 puncta and a different distribution of CTBP2 puncta compared to wildtype; 3) the *myo7aa^-/-^* mutants have significantly fewer numbers of active post-synaptic cells and fewer total MAGUK puncta; 4) the turning angles of the *myo7aa^-/-^* mutants are larger compared to wildtype turning angles; and 5) the *myo7aa^-/-^* mutants lack startle to a sound stimulus. We hypothesized that using L-type voltage gated calcium channel agonists will restore the abnormal phenotypes. We discovered that in the *myo7aa^-/-^* mutants 1) (R)-Baclofen increases ribbon area and number of tethered vesicles 2) All three L-type voltage-gated calcium channel agonists change the distribution of CTBP2 puncta to resemble wildtype distribution. 3) L-type voltage-gated calcium channel agonists improve swimming behavior and decrease turning angle 4) (±)-Bay K 8644 and Nefiracetam improve acoustic startle response in the *myo7aa^-/-^* mutants.

Ribbon synapses are only found in sensory cells of vertebrate animals. They are specialized presynaptic synapses that contain an electron dense structure that tethers synaptic vesicles near voltage-gated calcium channels at the active zone of the hair cell. The main component of a ribbon synapse is Ribeye with a monolayer of tethered glutamatergic synaptic vesicles (Schmitz et al., 2000). It has previously been reported that there are 3-5 single ribbons per hair cell in zebrafish (Nicolson, 2015). In this study we investigated if there is a difference in the ribbon synapse structure between the *myo7aa^-/-^* mutants and wildtype in ultrastructure and expression of genes involved in ribbon synapse structure and function. Through TEM we observed that synaptic ribbons were present at the synapse in the *myo7aa^-/-^* mutants indicating that the machinery to generate and localize the ribbons is functional in this model. We also observed that the ribbons in the *myo7aa^-/-^* mutants possess an electron dense structure that has a halo of tethered glutamatergic vesicles. Current understanding of this model’s hair cell is that it is not functional and is unable to transmit signal, however we challenge this view with our data that indicates the ribbons are generated with the correct components, are localizing to the synapse and the vesicles are fusing with the membrane to release neurotransmitter into the synaptic cleft. It has also been reported that hair cell innervation is important for the correct localization of the ribbons at the plasma membrane and for the regulation of ribbon size and number (Suli et al., 2016). They also conclude that the physical presence of afferent fibers is enough to regulate ribbon formation. Therefore, our observation of the presence of ribbon structures at the synapse in the *myo7aa^-/-^* mutants indicates that the hair cell is capable of stimulating afferent fibers. We argue that this signal in the *myo7aa^-/-^* mutants may not be sufficient to detect meaningful sound; however the ability of the cell to perform is intact. These observations support that cells are present and could respond to pharmacologic therapy. Additionally, Nicolson et al. conducted whole-cell patch-clamp recordings of wildtype and *myo7aa^-/-^* mutants and showed that the potential of the *myo7aa^-/-^* mutant is within the range of wildtype indicating that defects in this mutant hair cell do not cause an abnormal resting membrane potential and that the hair cell is still capable of transmitting a signal (Nicolson et al., 1998).

We identified that the area of the ribbon synapse does not differ between wildtype and *myo7aa^-/-^* mutants. It was also observed that ribbon size in another mechanotransduction mutant, *sputnik*-which harbors a mutation in *cadherin 23-* did not differ from wildtype (Suli et al., 2016). However, the *myo7aa^-/-^* mutants did have significantly fewer tethered vesicles. This further prompts the questions as to why there are fewer tethered vesicles. We hypothesize that this could be due to a decreased number of free cytosolic synaptic vesicles that would then be recruited at the opposite pole of the ribbon to replace docked vesicles that exocytose. This decrease in cytosolic synaptic vesicles would then lead to a decreased number of docked vesicles and therefore less neurotransmitter released into the synaptic cleft. In order to test this hypothesis cytosolic synaptic vesicles and docked vesicles would need to be quantified. Another possibility that could explain why there are fewer tethered vesicles in the *myo7aa^-/-^* mutants is that there is a disruption in the checkpoint for the attachment of synaptic vesicles to the dense body.

CTBP2 is alternatively spliced to produce CTBP2 and Ribeye, which is the central component of a ribbon synapse (Sheets et al., 2011, Wan et al., 2005). Sheets et al. identified that in zebrafish Ribeye is essential to cluster L-type voltage-gated calcium channels (Ca_v_1.3a) to the active zone at the basal end of the hair cell (2011). This was also described in cochlear inner ear hair cells (Frank et al., 2010). We explored the expression profile of CTBP2 in the *myo7aa^-/-^* mutants and discovered that *myo7aa^-/-^* mutants have significantly fewer active hair cells –defined as a hair cell containing at least one CTBP2 puncta-, fewer total CTBP2 puncta and a different distribution of CTBP2 puncta compared to wildtype. It was previously reported that hair cells of zebrafish larvae typically contain 3-5 ribbons (Nicolson, 2015, Obholzer et al., 2008). We identified that most wildtype hair cells contain three CTBP2 puncta and that most *myo7aa^-/-^* mutant hair cells have two CTBP2 puncta. Regus-Leidig et al. identified that regulation of RIBEYE A and B-domains controls the assembly and plasticity of synaptic ribbons and interactions between these two domains are inhibited by low concentrations of NAD(H^+^) (Magupalli et al., 2008, Regus-Leidig et al., 2010). Therefore, we suspect that the decreased expression of CTBP2 in the *myo7aa^-/-^* mutants could be explained by possible mitochondrial defects, however further analysis of mitochondria structure, function and by-products are necessary.

It is well established that pre-synaptic cell activity affects post-synaptic cell activity and vice versa (Luscher et al., 2000, Matus, 1999, Muller, 1997). Therefore, we used the post-synaptic cell marker, MAGUK to assess the post-synaptic cell and how it relates to what was observed in the ribbon synapse in the pre-synaptic cell. MAGUK is a kinase family that targets and anchors glutamate receptors to the synaptic terminal (Oliva et al., 2012). We discovered that *myo7aa^-/-^* mutants have significantly fewer number of active post-synaptic cells-defined as a cell with at least one MAGUK puncta- and fewer total MAGUK puncta per neuromast. We identified that most wildtype hair cells contain two to three MAGUK puncta and that most *myo7aa^-/-^* mutant hair cells have three MAGUK puncta. This is the first report of the distribution of CTBP2 and MAGUK puncta in wildtype and in any hearing defective mutant. By understanding how hair cell biology affects the post-synaptic cell we are able to speculate as to why the hair cell functions in a specific way. We hypothesize that the smaller number of CTBP2 puncta in the *myo7aa^-/-^* mutant is because the hair cell is not reaching the threshold required for normal expression of CTBP2. This may be due to a decrease in intracellular calcium concentrations or a decrease in expression of calcium binding proteins required for ribbon synapse formation. The decreased CTBP2 expression coupled with the data that the ribbon is localized to the synapse supports the hypothesis that the hair cell is releasing neurotransmitter into the synaptic cleft. It is interesting to note that although the total number of active post-synaptic cells and total number of MAGUK puncta is decreased, the distribution of the puncta is not different from wildtype. We know that MAGUKs are scaffolding proteins that maintain and modulate synaptic strength; therefore we believe that the fewer total number of MAGUK puncta in *myo7aa^-/-^* mutant post-synaptic cells is a compensatory mechanism in which the post-synaptic cells are responding to the decreased glutamate release from the hair cell. We can conclude that there is some regulation of AMPA and NMDA receptors leading to synaptic plasticity. This would need to be confirmed by analyzing the number of intact synapses by co-labeling with both antibodies and then determining if there are orphan post-synaptic densities.

A large-scale mutagenesis screen conducted in 1996 resulted in a large number of homozygous recessive mutants that exhibit behavioral defects (Granato et al., 1996). This study also included 15 morphologically normal mutants with balance abnormalities including the *mariner* mutant which displays looping and circular swimming starting at 4 dpf, develops a curved spine (that augments the abnormal swimming) at 5 dpf, and fails to develop a properly inflated swim bladder. We utilized a previously published behavior assessment that includes tracking zebrafish swimming in response to a static stimulus and were able to quantify this abnormal swimming behavior. We quantified the absolute smooth orientation, which is the sum of all turning angles a fish swam in one swimming episode as a function of time. The absolute smooth orientation of *myo7aa^-/-^* mutants is much larger compared to wildtype. This is also observed in the swimming trajectories where the *myo7aa^-/-^* mutants display circular and looping swimming and the wildtype animals have smooth trajectories and swim close to the edge of the well.

The objective of this study was to test the hypothesis that L-type voltage-gated calcium channel agonists can improve the abnormal cellular and behavioral phenotypes observed in the *myo7aa^-/-^* mutant. The rationale for this hypothesis stems from understanding that the *myo7aa^-/-^* mutant does not have a functional myo7aa protein therefore is unable to properly gate the MET channel, the membrane potential required for depolarization is not achieved, the L-type voltage-gated calcium channel does not open and synaptic transmission does not occur. We hypothesize that increasing the downstream signal in *myo7aa^-/-^* mutant hair cells by increasing intracellular calcium can reconstitute a new functional response to sound by increasing the sensitivity of the calcium channel through treatment with L-type voltage-gated calcium channel agonists. The three drugs we tested are (±)-Bay K 8644, Nefiracetam, and (R)-Baclofen. (±)-Bay K 8644 is a potent L-type calcium channel activator (Nowycky et al., 1985, Schramm et al., 1983). It has been used extensively to study regulation of calcium influx and the role of L-type calcium channels in synaptic transmission in neuronal cells (Bonci et al., 1998, Monjaraz et al., 2000, Watanabe et al., 1998). Nefiracetam is a nootropic or cognitive enhancer that activates both L and N-type calcium channels (Yoshii and Watabe, 1994, Yoshii et al., 2000) and may modulate the GABA receptor currents by interacting with a PKA pathway (Huang et al., 1996) and PKC pathway (Nishizaki et al., 1998). Nootropic agents can influence various neurotransmissions including dopaminergic, cholinergic, glutamatergic and GABAergic pathways (Funk and Schmidt, 1984, Giovannini et al., 1993, Huang et al., 1996, Marchi et al., 1990, Nabeshima et al., 1990, Nishizaki et al., 1998, Oyaizu and Narahashi, 1999, Watabe et al., 1993). Additionally, it has been demonstrated that Nefiracetam affects acetylcholine induced currents and NMDA receptors (Moriguchi et al., 2003, Narahashi et al., 2003, Zhao et al., 2001). NMDA and AMPA receptors are ionotropic glutamate receptors that are nonselective cation channels (K^+^ and Na^+^). Activation of these channels produces excitatory post-synaptic responses. NMDA receptors allow not only the entry of potassium and sodium ions, but also calcium; this increase in intracellular calcium acts as a secondary messenger to augment intracellular signaling cascades leading to the enrichment of synaptic activity. Previous studies have used Nefiracetam hypothesizing that this mechanism of action could improve post-stroke depression, post-stroke apathy and Alzheimer’s disease (Robert G. Robinson et al., 2009, Starkstein et al., 2016). Lastly, (R)-Baclofen was previously shown to act as GABA_B_ agonist (Bowery et al., 1983). GABA_B_ receptors are coupled through G-proteins to potassium channels, calcium channels, and adenylate-cyclase. However, more recently GABA_B_ receptors have been shown to modulate various voltage-gated calcium channels (Carter and Sabatini, 2004, Shen and Slaughter, 1999), specifically a direct interaction was identified between the L-type voltage-gated calcium channel (Ca_v_1.3) and GABA_B_ receptor subunit 2 (Park et al., 2010). They discovered that activation of the GABA_B_ receptor increases L-type calcium currents. (R)-Baclofen has been used in multiple recent clinical trials to treat Fragile X (Berry-Kravis et al., 2017) and autism (Veenstra-VanderWeele et al., 2017); and racemic Baclofen is an FDA approved drug currently used to treat muscle spasticity (Ertzgaard et al., 2017, Rizzo et al., 2004).

Modulating the L-type voltage-gated calcium channel activity was the central aim for using these three compounds. Sheets et al. reports that Ca_v_1.3a conductance affects the morphology of ribbon synapses; specifically identifying that 3 dpf zebrafish embryos had a decrease in Ca_v_1.3a conductance causing a larger ribbon area, but when they blocked Ca_v_1.3a with Isradipine at 5 dpf there was no change indicating that there is not much of an effect at more mature ribbon synapses (Sheets et al., 2012). They suggested that there is a period of synaptic maturation that is highly influenced by calcium flux through the Ca_v_1.3 that affects synaptic ribbons. We argue that this point is at 4 dpf in the development of inner ear hair cells and is the reason we chose to treat zebrafish at this developmental state for an overnight incubation.

Using TEM, we demonstrate that (R)-Baclofen increases ribbon area and number of tethered vesicles in the *myo7aa^-/-^* mutant ribbon synapse which is indistinguishable from wildtype. Incubation with (±)-Bay K 8644 and Nefiracetam did not have an effect on ribbon area and number of tethered vesicles on the *myo7aa^-/-^* mutants. Therefore, the mechanism in which (R)-Baclofen acts on the *myo7aa^-/-^* mutant ribbon synapses to increase ribbon area and number of tethered vesicles may not be directly and exclusively attributed to agonistic action of Ca_v_1.3a, however our study is the first to demonstrate that ribbon synapses of the *myo7aa^-/-^* mutant can be altered with drug incubation to more closely resemble wildtype ribbon synapses. We did not see any adverse effects in the ribbon area or number of tethered vesicles on wildtype larvae upon incubation with the three drugs.

In addition to demonstrating that (R)-Baclofen can alter ribbon synapse area and number of tethered vesicles, we demonstrated that all three compounds change the distribution of CTBP2 puncta. Our finding that most wildtype hair cells have two CTBP2 puncta and most *myo7aa^-/-^* mutant hair cells have three CTBP2 puncta establishes that this difference may be contributing to the phenotype of the *myo7aa^-/-^* mutant. Our data shows that incubation with all three L-type voltage-gated calcium channel agonists shifts the distribution of CTBP2 puncta and restores the distribution to resemble wildtype. This is an important discovery, because it indicates that all three compounds are likely acting in the same manner in their effect on CTBP2 distribution. Since (±)-Bay K 8644 only has one mechanism of action, L-type voltage-gated calcium channel agonist-we can presume that Nefiracetam and (R)-Baclofen effects may be primarily through the same pathway. Increasing intracellular calcium through this mechanism may have an effect on the regulation of CTBP2. We also identified that (±)-Bay K 8644 significantly increased the total number of active hair cells and post-synaptic cells per neuromast and the total number of CTBP2 and MAGUK puncta per neuromast. It is likely that only (±)-Bay K 8644 had this effect because it is the most potent of the three and its only mechanism of action is to act on the Ca_v_1.3a channel. We also observed that upon incubation with 5 μM (±)-Bay K 8644 the distribution of CTBP2 puncta did shift in wildtype larvae so that most hair cells have two CTBP2 puncta rather than three which is observed in the untreated wildtype groups. One possible explanation could be that we are observing a dose response that takes a bell-shaped curve meaning that some doses (lower or higher) do not give a specific outcome whereas those in the middle do. In this case, we suspect that 5 μM is a dose that falls in the middle of a bell shaped curve and would elicit a response. This hypothesis could be tested by measuring CTBP2 expression with higher and lower doses of (±)-Bay K 8644. Recent work shows that evoked presynaptic calcium influx through L-type voltage-gated calcium channels stimulates calcium uptake in the mitochondria causing decreased cellular NAD(H) thus decreasing ribbon formation (Wong et al., 2019), which may explain why 5 μM (±)-Bay K 8644 shifts the distribution of CTBP2 puncta in wildtype hair cells, however this is unlikely in this situation because we see that (±)-Bay K 8644 actually increases the total number of active hair cells and post-synaptic cells per neuromast and the total number of CTBP2 and MAGUK puncta per neuromast. Lastly, (R)-Baclofen also increased the number of active post-synaptic cells and total number of MAGUK puncta in wildtype larvae.

It is well characterized that zebrafish demonstrate a short-latency startle response, the C-start, in response to tactile and auditory stimuli that consists of a C-bend of the body followed by a counter bend and then a swimming burst (Eaton et al., 1991, Kimmel et al., 1974). This response is a result of activation of the Mauthner cells (M-cells). Zottoli et al. linked Mauthner cell activity to escape behavior (Zottoli, 1977). We utilized a behavior assessment that tracks zebrafish swimming behavior and quantifies the absolute smooth orientation or turning angle as a function of time in response to a static stimulus. We identified that the absolute smooth orientation of *myo7aa^-/-^* mutants is much larger compared to wildtype. We showed improved swimming behavior in the *myo7aa^-/-^* mutants meaning a decrease in turning angle using all three L-type voltage-gated calcium channel agonists; however the most robust response was with (±)-Bay K 8644, which resulted in turning angles indistinguishablefrom wildtype. We tested all three drugs in wildtype animals as well and observed no change in the absolute smooth orientation at these specific doses with the exception of (±)-Bay K 8644. This could be explained by the fact that the doses chosen for Nefiracetam and (R)-Baclofen were chosen based on dose-response studies (Figs. S4 & S6). These data show that changes in the absolute smooth orientation are observed in wildtype larvae depending on the dose. We did not conduct a dose response study for (±)-Bay K 8644, therefore we would expect that other doses would result in no significant changes in swimming behavior.

Not only did our quantitative analysis of the swimming behavior indicate improvement, swimming trajectories of individual fish from each group also demonstrate less circular and looping swimming and more smooth and zig-zag swimming. It is interesting to note that all wildtype fish (untreated or treated) had a preference for swimming toward the edge of the well, a process called thigmotaxis. Peitsaro et al. report that thigmotaxis in zebrafish is an indicator of anxiety (Peitsaro et al., 2003). This has also been shown in mice (Simon et al., 1994). It is unclear if this process is disrupted in the *myo7aa^-/-^* mutants or if the abnormal swimming behavior is what prevents the fish from remaining at the edge of the well.

The acoustic startle response can be elicited starting at 5 dpf, which is when the ear is completely developed (Eaton and Didomenico, 1986, Zeddies and Fay, 2005). The zebrafish has two sensory organs: the inner ear and lateral line system. The inner ear contains three otolithic organs: the utricle, the saccule and at 11-12 dpf the lagena develops (Popper and Fay, 1993). Saccular hair cells respond to relatively high frequencies (200 Hz-1200 Hz) and fibers that innervate the saccule synapse on M-cells (Bang et al., 2002, Fay, 1995, Lin and Faber, 1988, Zottoli, 1977). The lateral line responds to much lower frequencies (50-100 Hz) (Coombs, 1994, Coombs and Janssen, 1990). There are multiple hearing assessments previously reported including acoustically evoked behavioral responses, tone burst, pre-pulse inhibition assays, microphonic potentials and auditory brainstem responses (Bang et al., 2002, Bhandiwad et al., 2013, Burgess and Granato, 2007, Higgs et al., 2002, Yang et al., 2017, Yao et al., 2016, Zeddies and Fay, 2005). However, all of these assessments require specialized equipment and analysis tools that would provide more information than what we required to asses presence or absence of an acoustic startle response. We, therefore, sought to develop a cost-effective and accurate hearing assessment tool that can be accessible to others. In our hearing assessment we acquired videos of 5 dpf larval fish placed on a speaker and administered frequencies ranging from 200 to 600 Hz. We used *myo7aa^-/-^* mutants as a control for the experimental design and utilized an intensity level that stimulated the wildtype fish but did not cause a startle in the *myo7aa^-/-^* mutants. Using this paradigm, we identified that untreated wildtype fish do not respond with a 100% response rate at any frequency and we see the most robust responses at 200 and 400 Hz. Incubation with (±)-Bay K 8644 and (R)-Baclofen decreased mean response rate in the wildtype fish, however incubation with Nefiracetam improved the mean response rate. A potential explanation as to why we are observing adverse effects in wildtype fish upon incubation with (±)-Bay K 8644 in not only the acoustic startle but also in the distribution of CTBP2 puncta and the absolute smooth orientation is that excessive cytosolic calcium (altering calcium homeostasis) in the hair cell can be detrimental (Esterberg et al., 2013). Another explanation could be that the elevated cytosolic calcium in the wildtype hair cells due to (±)-Bay K 8644 could be causing excessive activation of the glutamate receptors that has been linked to apoptotic hair-cell death (Sheets, 2017). We may be observing this affect across various assessments with only (±)-Bay K 8644, in the wildtype animals because it is the more potent drug. In other words, in a normal hair cell if the calcium concentration falls above the threshold of homeostasis then it can have adverse effects on the cell. However, in a hair cell that already has a starting calcium concentration below the threshold (*myo7aa^-/-^* mutant) than pharmacologic strategies to increase intracellular calcium will bring the calcium concentration to the level of homeostasis.

It has previously been reported that incubation with (R)-Baclofen can cause a sedative effect in zebrafish that decreases locomotor activity that is dose dependent (Renier et al., 2007). This study reports that doses as low as 10^-4^ M dropped the locomotor activity to about 25-30% of what was observed in the controls. We are seeing that wildtype fish treated with 125 µM (R)-Baclofen exhibit about 20% of the mean response rate of control fish at each individual frequency.

Lastly, we tested the hypothesis that L-type voltage gated calcium channel agonists can restore some hearing by increasing mean acoustic startle response rate in the *myo7aa^-/-^* mutants. Our data suggests that (±)-Bay K 8644 and Nefiracetam improve acoustic startle response in the *myo7aa^-/-^* mutants. This is an important discovery because it is the first report of a pharmacotherapy restoring hearing in a zebrafish model of USH1. Additionally, like other otophysan fishes, the zebrafish has a swimbladder that is coupled to the saccule through Weberian ossicles that facilitates responses to sound pressure (Higgs et al., 2003). The *myo7aa^-/-^* mutant does not develop a swimbladder. We tested if L-type voltage gated calcium channels affect inflation of the swim bladder in the *myo7aa^-/-^* mutant and identified that Nefiracetam and (R)-Baclofen did not result in an inflated swim bladder and did not negatively affect wildtype swim bladders. This supports our hearing assessment data suggesting that the *myo7aa^-/-^* mutant fish incubated in L-type voltage-gated calcium channel agonists that are responding to acoustic stimuli are responding to direct acceleration of the cells within the otolith organs, specifically the saccule and not to changes in sound pressure.

## Conclusion

Here we report the first study using L-type voltage-gated calcium channel agonists to treat the zebrafish model of USH1, the *myo7aa^-/-^* mutant. We have shown through transmission electron microscopy and immunohistochemistry that the hair cell biology of the *myo7aa^-/-^* mutant differs from wildtype in that there are fewer glutamatergic vesicles tethered to the ribbon synapse, fewer total active hair cells and post-synaptic cells, fewer total CTBP2 and MAGUK puncta, and a different distribution of CTBP2 puncta. We quantified the previously observed behavioral phenotypes and identified that *myo7aa^-/-^* fish have larger turning angles and little to no acoustic startle response. Upon treatment with L-type voltage-gated calcium channel agonists, we discovered changes in hair cell morphology/ultrastructure, gene expression and behavior through swimming and hearing assessments. Our data support that treatment with these compounds restores the abnormalities in the *myo7aa^-/-^* mutants to more closely resemble wildtype by 1) increasing the number of tethered vesicles to the ribbon 2) shifting the distribution of CTBP2 puncta 3) decreasing turning angle and improving swimming behavior and 4) improving acoustic startle. However, there are still many unanswered questions in regards to localization and density of L-type voltage-gated calcium channels, ratio of ribbons at the synapse versus those that are not, mitochondrial contribution to the phenotype or the rescue and direct quantification of calcium concentrations with and without treatment. We are confident that the results we report in this study represent a significant step towards understanding the mechanism of disease and discovering compounds to treat hearing loss caused by pathogenic variants in *MYO7A*.

## MATERIALS AND METHODS

### Zebrafish maintenance and husbandry

All animals used in this study were zebrafish (*Danio rerio).* The zebrafish strains used for all experiments were the tc320b allele (c.2699T>A; p.Tyr846Stop) in exon 21 of *myo7aa* (Ernest et al., 2000) and wildtype Segrest strains were obtained from the laboratory stock at Mayo Clinic (Leveque R.E, 2016). Heterozygous adults for the *myo7aa* mutation were used to generate homozygous larval fish (*myo7aa^-/-^*). Animals were raised in a 10-hour dark and 14-hour light cycle with embryos and adults kept at 28.5°C (Westerfield, 1995). Embryos were kept in Petri dishes and fresh embryo water was replaced daily. Larvae were kept at room temperature for all experiments and were maintained in the incubator at all other times. Wildtype Segrest strains and heterozygous adult *myo7aa* fish were also maintained at the University of Minnesota, Twin Cities Zebrafish Core Facility. The Institutional Animal Care and Use Committee approved all experimental procedures both at Mayo Clinic and the University of Minnesota.

### Phenotyping and genotyping

The *myo7aa^-/-^* tc320b allele results from a single nucleotide polymorphism A-T, which results in an early stop codon p.Tyr846Stop (Ernest et al., 2000). Homozygous recessive mutant larval fish were identified at 4 dpf (days post fertilization) based on various phenotypic traits, including defective balance, absent startle to sound, and circular or looping swimming patterns. DNA from whole embryos or fin biopsies was extracted for allele-specific quantitative PCR for genotyping as previously described (Lee et al., 2016).

### Hair cell staining

In order to stain the stereocilia of the saccule hair cells, embryos were initially fixed overnight in freshly thawed 4% paraformaldehyde at 4°C. Two washes for 5 minutes each in 0.2% PBSTx were conducted at room temperature. Embryos were then incubated for 4 days in 2% PBSTx at 4°C to completely dissolve the otoliths. F-actin staining occurred for up to 2 hours at room temperature using Alexa Fluor 488 Phalloidin (Molecular Probes, Eugene, OR), which was diluted 1:20 in PBSTw. Embryos were then washed six times for 10 minutes at room temperature in 0.2% PBSTx. Hair cell stereocilia were observed and imaged using the LSM 780 inverted confocal microscope run on Zen software package (Zeiss).

### MET channel assessment

Mechanotransduction channel activity was assessed through FM 1-43 dye uptake (Molecular Probes, Eugene, OR). A working stock of 3 µM FM 1-43 was used for a 30 second incubation and rinsed three times immediately with embryo water. Lateral line neuromasts were imaged for FM 1-43 uptake using a Lightsheet Z.1 microscope (Zeiss). Lateral-orientated z-stacks at 5X/1.0 NA water-dipping objective (Zeiss) and an RFP (emission filter BP 505-545 LP 585) optical filter (Zeiss). Each corresponding dorsal image is a maximal image projection generated from z-dimension stacks.

### Drug incubation

The maximum tolerated dose was determined for all drugs used in this study. This was performed by incubating 4 dpf wildtype embryos individually in 24-well plates in 1 mL total volume of embryo water including various doses of dissolved drug. Control fish were incubated in the same concentration of embryo water with only drug vehicle (e.g. DMSO, H_2_O). The 24-well plates were placed in a 29°C embryo incubator for the duration of the drug treatment. Heart rates, startle responses, swimming behavior and overall health were observed after one hour, four hours and overnight incubation to determine any adverse effects or possible toxicity. Drug doses started at 0.5 μM and increased based on overall health and survival of the embryos. The drugs used in this study are as follows: (±)-Bay K 8644 (catalog # 1544, Batch No. 4; Tocris), Nefiracetam (catalog # 2851, Batch No. 1; Tocris) and (R)-Baclofen (catalog # 0796, Batch No. 3; Tocris). (±)-Bay K 8644 and Nefiracetam were dissolved in DMSO to a 100 mM stock solution and (R)-Baclofen was dissolved in water (with gentle warming) to a 20 mM stock solution. The final doses that would be used in the drug treatments were determined by identifying the highest tolerated dose and using the dose one step lower. A control experiment was conducted to determine the effect of DMSO on behavior and at the concentrations used for dissolving drugs, and no adverse effects were identified.

### Video acquisition of swimming behavior

Videos were made according to the video acquisition protocol previously described (Lambert et al., 2012). High-speed videos of 5dpf larvae were acquired at either 120 or 180 frames per second (fps) using FlyCap software and the Point Grey Firefly. A LED light stage was used underneath a 20 mm arena in order to increase illumination for a higher quality video. A 50 V electric shock (5 ms duration) was used to startle the animal every 20 seconds. An LED light was used to confirm the administration of the electric shock.

### Video tracking and analysis

Videos acquired from the FlyCap software were processed as fmf files through Ctrax: The Caltech Multiple Fly Tracker, free source software (http://ctrax.sourceforge.net/) (Branson et al., 2009). The tracking settings for motion, observation and hindsight are listed in Table S1. Videos were exported as mat files, fixed for errors using the Fix Errors Matlab Toolbox (FEMT), provided by (Branson et al., 2009), and analyzed in Matlab version 2012b using in-house scripts from the Masino Laboratory (available upon request). Swimming trajectories were generated using the Ctrax Behavioral Microarray Matlab toolbox. A circle indicating the perimeter of the testing well was added to each image as a reference point. Each electric shock was confirmed by determining the frame in which the LED light appears using Image J. The frame range corresponding to that swimming episode was found in the mat files. The values for the absolute smooth orientation of the fish during one swimming episode were totaled and then divided by the episode duration. This was performed for all swimming episodes.

### Hearing assessment

Up to ten 4 dpf embryos were placed in 60 × 15 mm Petri dishes and placed in a 29°C embryo incubator in an isolated behavior room for overnight incubation. At 5dpf embryos were placed on top of a Bose Soundlink Color Speaker and allowed up to 5 min to acclimate. The speaker was placed on top of an illuminator for a better quality video. A white piece of paper was placed beneath the Petri dish to lighten the background in order to create contrast between the larval fish and the speaker. A script written in Matlab (available upon request) was used to generate 3 one second stimuli every 20 seconds for the following frequencies: 200, 300, 400, 500 and 600 Hz and administered via Bluetooth connection between the laptop and the speaker. The intensity of the sounds administered was determined using an Extech sound level meter, around 77 dB. Videos were taken using a Sony Handycam HDR cx560 and were de-identified and blindly scored by a trained laboratory member for startle responses. Responses were scored as either 0 (no response) or 1 (response) for each fish at each stimulus. Mean values of responses and no responses were calculated and the mean variance was determined.

### Processing and imaging of zebrafish for TEM

5dpf embryos were fixed in Trump’s fixative (4 % paraformaldehyde with 1% glutaraldehyde) for a minimum of 24 hr. Following fixation, embryos were secondarily fixed with 1% osmium tetroxide and 2% uranyl acetate, dehydrated through an ethanol series and embedded into Embed 812/Araldite resin. Processing was facilitated with the use of a BioWave laboratory microwave oven (Pelco Biowave 3450, Ted Pella, Inc., Redding, CA). After a 24 hr polymerization at 60°C, ultrathin sections (0.1 mm) were stained with 2% uranyl acetate and lead citrate. Micrographs were acquired in the Mayo Clinic Microscopy and Cell Analysis Core using a JEOL1400 or JEOL1400Plus transmission electron microscope operating at 80 kV (Peabody, MA) equipped with a Gatan Orius camera (Gatan, Inc., Warrendale, PA).

### Analysis of ribbon synapse TEM images

Transmission electron microscopy images of ribbon synapse structures were de-identified and analyzed for ribbon area and number of tethered vesicles. ImageJ software was used to determine the area of the electron dense structure and the number of vesicles was quantified manually.

### Immunohistochemistry

The preparation and the immunohistochemistry (IHC) experiments were conducted across five days using a nutator unless otherwise specified. The first day included fixing 5dpf embryos in 4% paraformaldehyde at 4°C for overnight incubation. Samples were then rinsed twice and washed three times for 10 minutes each with PBTr (0.02% Triton-X 100 in 1X PBS), washed for 5 minutes in a 1:1 mixture (PBTr and 100% MetOH) and then washed three times for 5 minutes in 100% MetOH. They were stored at −20°C if not used immediately. The third day included rehydrating the embryos through a 5-minute wash of the 1:1 mixture followed by three 5-minute washes with PBTr. Samples were permeabilized for 50 minutes with proteinase K (diluted 1:2000 in PBTr) and then washed five times for 5 minutes with PBTr. Block buffer included 10% Normal Goat Serum, 0.2% Triton-x 100, 2% DMSO in 1X PBS. Samples were incubated in block buffer for at least 3 hours. The primary antibodies used in this study were mouse monoclonal against membrane associated guanylate kinase, a post-synaptic membrane marker (MAGUK; 1:250 NeuroMab Clone K28/86, Catalog # 75-029) (Anderson 1996; Lv et al. 2016; Sheets et al. 2011)) and CTBP2 (1:500 Santa Cruz Catalog # Sc-55502)(Lv et al. 2016) diluted in prepared block buffer. Samples were incubated in diluted primary antibody at 4°C overnight. The fourth day included washing samples six times for 30 minutes with PBTr. The secondary antibodies used were Alexa Fluor 647 goat anti-mouse (1:250 Thermo-Fisher/Invitrogen Catalog # A21235) and Alexa Fluor 488 goat anti-mouse (1:500 Thermo-Fisher/Invitrogen Catalog # A21121) respectively and were diluted in PBTr. Samples were incubated in diluted secondary antibody at 4°C overnight. Lastly, samples were washed again six times for 30 minutes each wash with PBTr and washed three times for 5 minutes with 1X PBS. Samples were stored in 1X PBS in 4°C. Images were acquired using the LSM 780 inverted confocal microscope run on Zen software package (Zeiss). Samples were mounted with Vectashield Hardset DAPI Mounting Medium, and covered with a cover slip for approximately 10 minutes before imaging. All images were captured with the 20x objective and a 5.0 magnitude zoom. All images were de-identified prior to analysis using ImageJ software. The proportion of active cells with 1,2,3,4,5,6+ puncta was determined by identifying the number of cells with n puncta divided by the total number of active cells in that specific neuromast.

### Statistical analysis

Statistical analyses were performed using JMP software (JMP^®^ 1989-2019). Each treatment group was individually compared to the corresponding control group (wildtype or *myo7aa^-/-^* untreated) through a student’s t-test. Acoustic startle response and CTBP2/MAGUK puncta statistical analysis was performed using a two-tailed Fischer’s exact test (https://www.graphpad.com/quickcalcs/contingency1/). All statistical analyses are presented with appropriate significant digits.

## Supporting information

Supplemental Data Koleilat et al

## Acknowledgements

We thank Dr. Teresa Nicolson for providing us with the tc320b allele of *mariner*. We are grateful to Mark Curry and Patrick Jochim in the Mayo Clinic Media Services. We also thank Gabriel Martinez and Dr. Timothy Wiggins for help with Adobe Illustrator, Matlab and the behavior assessment. We are also grateful to Dr. Lavinia Sheets for her helpful discussions. Lastly, we would like to thank the staff members in the zebrafish core facilities at Mayo Clinic and the University of Minnesota, Twin Cities.

## Data Availability

All data required for analysis is included within the main text.

**Author Contributions**

Conceptualization-A.K, S.C.E, L.A.S

Methodology-A.K, A.M.L, M.A.M

Software-A.K, A.M.L, M.A.M

Validation-A.K, J.D, M.A.M

Formal Analysis-A.K

Investigation-A.K, J.D, J.B, T.C

Resources-A.K, J.D, M.A.M, S.C.E, L.A.S

Data Curation-A.K., A.M.L, M.A.M

Writing-original draft preparation-A.K.

Writing-review and editing-A.K, J.D., T.C., J.B., A.M.L, M.A.M, S.C.E, L.A.S

Visualization-A.K.

Supervision-S.C.E, L.A.S

Project administration-A.K, S.C.E, L.A.S

Funding acquisition-L.A.S

