## Supplemental Data Koleilat et al for "L-type voltage-gated calcium channel agonists improve hearing loss and modify ribbon synapse morphology in the zebrafish model of Usher Syndrome Type 1"

| Setting | Value |
| --- | --- |
| Angle Weight | 100 |
| Max Jump Distance | 80 |
| Max Jump D to Split | 80 |
| Min Jump Distance | 10000 |
| Center Dampening | 0 |
| Angle Dampening | 0.5 |
| Max Area Delete | 200 |
| Min Area Ignore | 20000 |
| Max Penalty Merge | 5000 |
| Lower Thresh | 1 |
| Max Clusters Per Blob | 1 |
| Max Blobs to Detect | 2 |
| Max Sequence Length | 120 or 180 (frame rate) |
| Max Penalty Merge | 5000 |
| Max Pred Error Increase | 5000 |

**Table S1. Tracking settings for motion, observation, and hindsight in Ctrax: The Caltech Multiple Fly Tracker.**

| | wildtype untreated | | wildtype untreated |
|---|---|---|---|
| CTBP2 Puncta | *myo7aa*$^{-/-}$ untreated | MAGUK Puncta | *myo7aa*$^{-/-}$ untreated |
| 1 | 1.0 | 1 | 0.36 |
| 2 | **0.0001** | 2 | 0.88 |
| 3 | **0.0003** | 3 | 0.15 |
| 4 | 0.33 | 4 | 0.06 |
| 5 | 0.15 | 5 | 0.70 |
| 6+ | 0.12 | 6+ | 0.31 |

**Table S2. *myo7aa*$^{-/-}$ mutant hair cells have a statistically different distribution of CTBP2 puncta, but a similar distribution of MAGUK puncta.** A two-tailed Fisher's exact test was used to determine statistical significance between wildtype and the *myo7aa*$^{-/-}$ mutant CTBP2 and MAGUK puncta. P-values are represented in each cell.

| A | wildtype untreated | | |
|---|---|---|---|
| CTBP2 Puncta | wildtype 5 µM (±)-Bay K 8644 | wildtype 250 µM Nefiracetam | wildtype 125 µM (R)-Baclofen |
| 1 | **0.0001** | 1.0 | 0.1651 |
| 2 | **0.0003** | 0.17 | **0.003** |
| 3 | **0.0005** | 0.17 | 0.31 |
| 4 | **0.002** | 0.55 | 0.72 |
| 5 | 0.06 | 1.0 | 0.09 |
| 6+ | 0.09 | 0.33 | **0.007** |

| B | *myo7aa*$^{-/-}$ untreated | | |
|---|---|---|---|
| CTBP2 Puncta | *myo7aa*$^{-/-}$ 5 µM (±)-Bay K 8644 | *myo7aa*$^{-/-}$ 250 µM Nefiracetam | *myo7aa*$^{-/-}$ 125 µM (R)-Baclofen |
| 1 | **0.04** | 0.25 | **0.01** |
| 2 | **0.002** | **0.01** | **0.02** |
| 3 | **0.008** | **0.003** | **0.03** |
| 4 | 0.65 | 1.0 | 1.0 |
| 5 | **0.02** | 0.06 | **0.005** |
| 6+ | 1.0 | 1.0 | 0.63 |

**Table S3. L-type voltage-gated calcium channel agonists change the distribution of CTBP2 puncta in *myo7aa*$^{-/-}$ mutant hair cells.** A two-tailed Fisher's exact test was used to determine statistical significance between treated and untreated groups. P-values are represented in each cell. Table A represents statistical analysis for all wildtype treated animals compared to untreated wildtype animals for each number of puncta. (±)-Bay K 8644 had the largest effect on wildtype hair cells. Table B represents statistical analysis for all *myo7aa*$^{-/-}$ treated animals compared to untreated *myo7aa*$^{-/-}$ animals for each number of puncta. All three L-type voltage-gated calcium channel agonists decreased the number of hair cells with 2 CTBP2 puncta and increased the number of hair cells with 3 CTBP2 puncta.

| A | wildtype untreated | | |
|---|---|---|---|
| MAGUK Puncta | wildtype 5 µM (±)-Bay K 8644 | wildtype 250 µM Nefiracetam | wildtype 125 µM (R)-Baclofen |
| 1 | 0.24 | 0.06 | 0.64 |
| 2 | 0.05 | **0.04** | 0.11 |
| 3 | 0.88 | 0.80 | 0.13 |
| 4 | **0.005** | 0.69 | **0.01** |
| 5 | 0.34 | 0.73 | 0.35 |
| 6+ | 0.59 | 1.0 | 0.14 |

| B | ***myo7aa***$^{-/-}$ **untreated** | | |
|---|---|---|---|
| MAGUK Puncta | *myo7aa*$^{-/-}$ 5 µM (±)-Bay K 8644 | *myo7aa*$^{-/-}$ 250 µM Nefiracetam | *myo7aa*$^{-/-}$ 125 µM (R)-Baclofen |
| 1 | 0.92 | 1.0 | 0.31 |
| 2 | 0.17 | **0.0001** | 0.46 |
| 3 | 0.81 | 0.36 | 0.21 |
| 4 | 0.56 | 0.51 | 0.44 |
| 5 | 0.65 | 0.88 | 0.22 |
| 6+ | 0.46 | 0.86 | 1.0 |

**Table S4. Overall, L-type voltage-gated calcium channel agonists do not change the distribution of MAGUK puncta in *myo7aa*$^{-/-}$ mutant hair cells.** A two-tailed Fisher's exact test was used to determine statistical significance between treated and untreated groups. P-values are represented in each cell. Table A represents statistical analysis for all wildtype treated animals compared to untreated wildtype animals for each number of puncta. Overall there was no change on the distribution of MAGUK puncta. Table B represents statistical analysis for all *myo7aa*$^{-/-}$ treated animals compared to untreated *myo7aa*$^{-/-}$ animals for each number of puncta. Overall, all three L-type voltage-gated calcium channel agonists had no effect on the number of MAGUK puncta in the post-synaptic cells.

| A | wildtype untreated | | |
|---|---|---|---|
| Frequency (Hz) | wildtype 5 μM (±)-Bay K 8644 | wildtype 250 μM Nefiracetam | wildtype 125 μM (R)-Baclofen |
| 200 | 0.34 | **0.0001** | **<0.0001** |
| 300 | **0.0009** | **0.003** | **<0.0001** |
| 400 | **0.04** | **0.01** | **<0.0001** |
| 500 | **<0.0001** | **0.03** | **<0.0001** |
| 600 | **0.002** | **0.006** | **0.0004** |

| B | *myo7aa*$^{-/-}$ untreated | | |
|---|---|---|---|
| Frequency (Hz) | *myo7aa*$^{-/-}$ 5 μM (±)-Bay K 8644 | *myo7aa*$^{-/-}$ 250 μM Nefiracetam | *myo7aa*$^{-/-}$ 125 μM (R)-Baclofen |
| 200 | **<0.0001** | **0.016** | 1.0 |
| 300 | **0.006** | 1.0 | 1.0 |
| 400 | **<0.0001** | **0.008** | 1.0 |
| 500 | **0.006** | 0.08 | 1.0 |
| 600 | **<0.0001** | 0.14 | 0.51 |

**Table S5. Statistical Analysis for hearing assessment per frequency.** A two-tailed Fisher's exact test was used to determine statistical significance between untreated and treated group. P-values are represented in each cell. Table A represents statistical analysis for all wildtype treated animals compared to untreated wildtype animals for each frequency. Table B represents statistical analysis for all *myo7aa*$^{-/-}$ mutants treated compared to untreated *myo7aa*$^{-/-}$ mutants for each frequency.

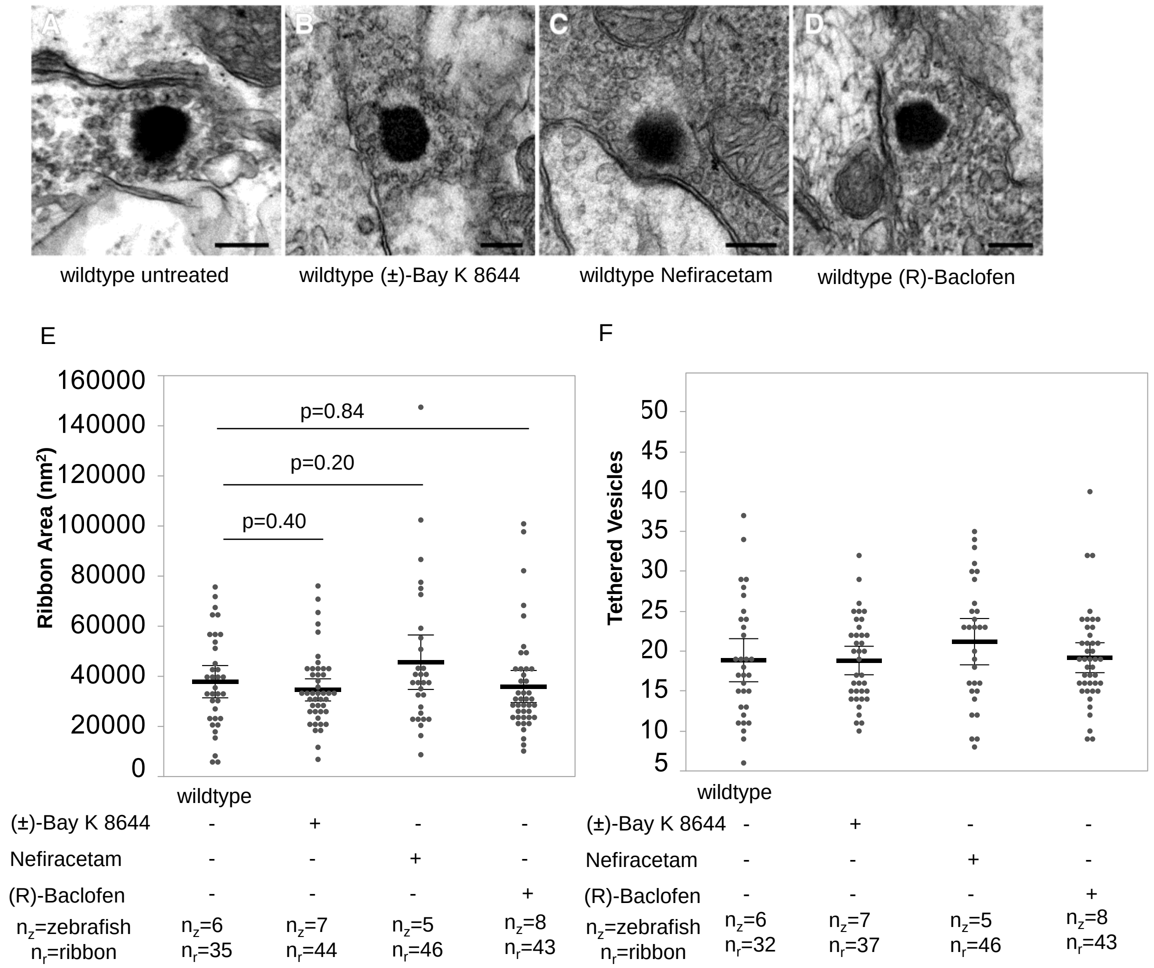


**Fig. S1. Wildtype ribbon synapses unaffected with exposure to L-type voltage-gated calcium channel agonists.** (A-D) Representative images of 5dpf wildtype untreated and treated with 5 µM (±)-Bay K 8644 (B), 250 µM Nefiracetam (C) and 125 µM (R)-Baclofen (D) ribbon synapse structure obtained with transmission electron microscopy. Scale bars: 200 nm. (E) Wildtype treated and untreated ribbon synapse areas are comparable. (F) Additionally, untreated and treated wildtype ribbons have a comparable number of tethered vesicles. Black bold line represents the mean of the data set and error bars are 95% confidence intervals.

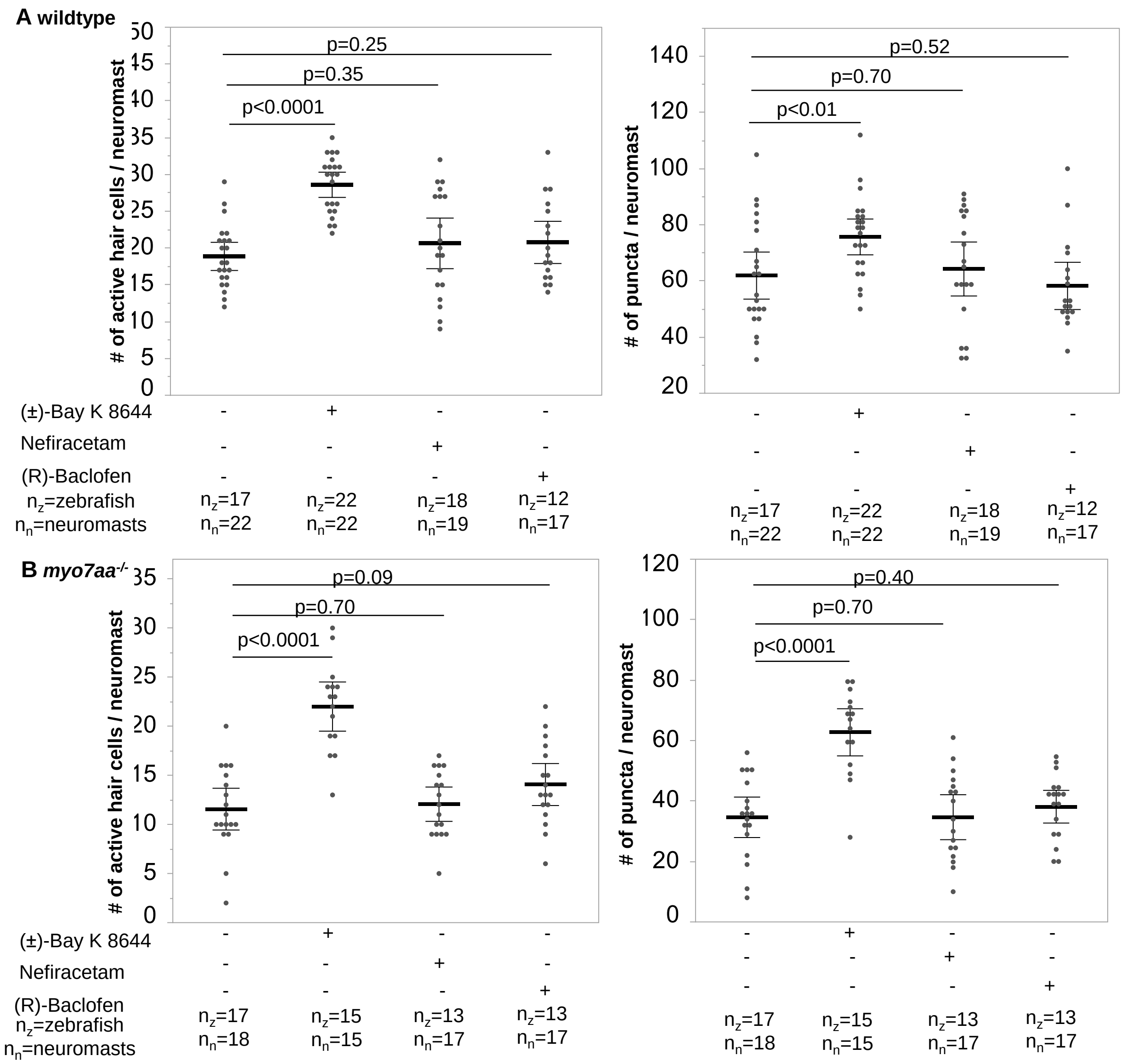


**Fig. S2 (±)-Bay K 8644 increases total number of active hair cells/neuromast and total number of CTBP2 puncta/neuromast in both wildtype and *myo7aa*$^{-/-}$ mutants.** (A) 5dpf wildtype neuromasts treated with (±)-Bay K 8644 have a greater number of active hair cells and total number of CTBP2 puncta compared to untreated wildtype neuromasts. An active hair cell is defined as a hair cell with at least one CTBP2 puncta (t-test). (B) 5dpf *myo7aa*$^{-/-}$ neuromasts treated with (±)-Bay K 8644 have a greater number of active hair cells and total number of CTBP2 puncta compared to untreated *myo7aa*$^{-/-}$ neuromasts (t-test). Black bold line represents the mean of the data set and error bars are 95% confidence intervals.

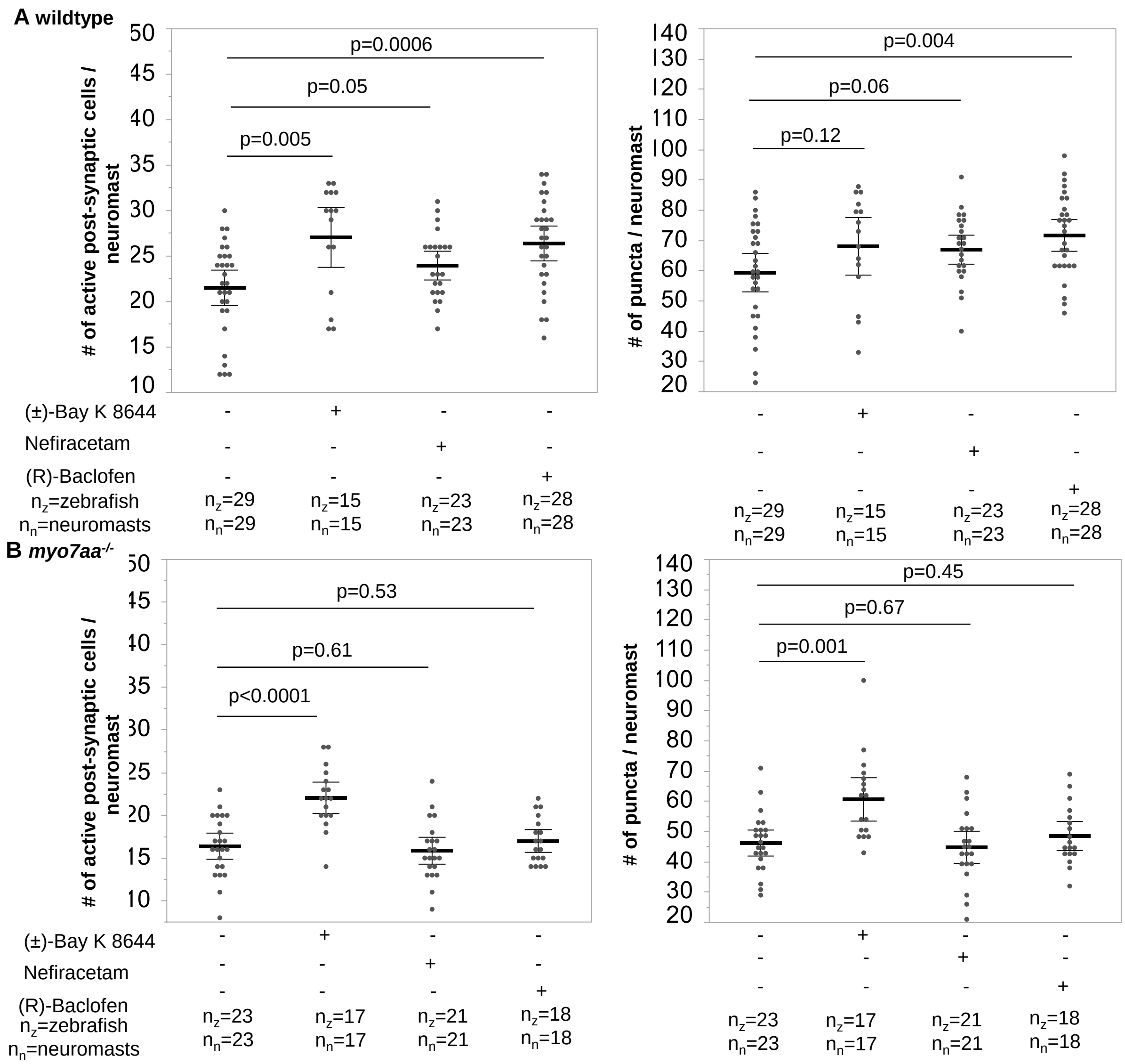


**Fig. S3 (±)-Bay K 8644 and (R)-Baclofen increase total number of active post-synaptic cells/neuromast and total number of MAGUK puncta/neuromast in wildtype however only (±)-Bay K 8644 affects *myo7aa$^{-/-}$* mutants.** (A) 5dpf wildtype neuromasts treated with (±)-Bay K 8644 and (R)-Baclofen have a greater number of active post-synaptic cells and total number of MAGUK puncta compared to untreated wildtype neuromasts. An active post-synaptic cell is defined as a cell with at least one MAGUK puncta (t-test). (B) 5dpf *myo7aa$^{-/-}$* neuromasts treated with (±)-Bay K 8644 have a greater number of active post-synaptic cells and total number of MAGUK puncta compared to untreated *myo7aa$^{-/-}$* neuromasts (t-test). Black bold line represents the mean of the data set and error bars are 95% confidence intervals.

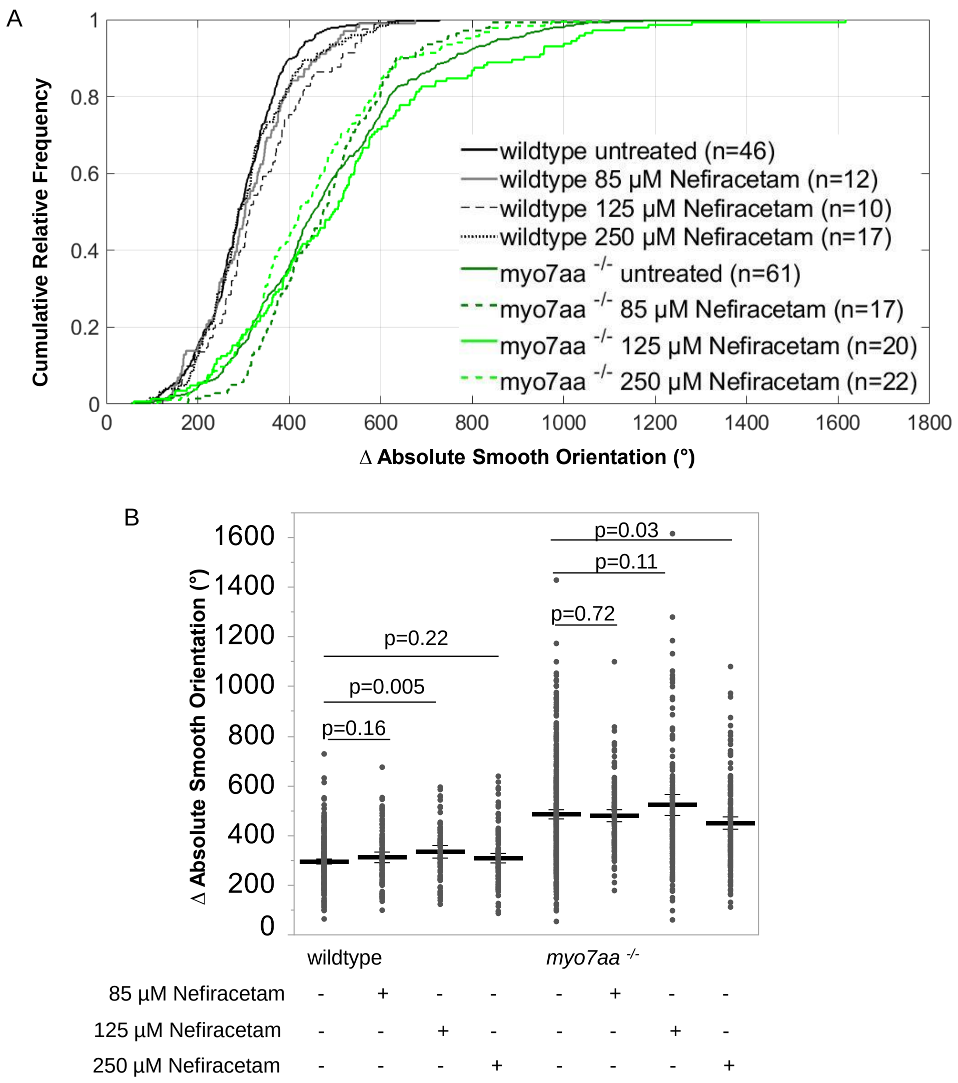


**Fig. S4 Different doses of Nefiracetam have different effects on wildtype and *myo7aa*$^{-/-}$ mutant swimming behavior.** This behavior assessment tracks movement of 5dpf wildtype and *myo7aa*$^{-/-}$ larvae over a 2.5 minute interval with a 5 ms electric stimulus (50 mV) administered every 20 s. Ctrax software was used for video processing and Matlab 2012b for video analysis. (A, B) Incubation with 85 µM and 250 µM Nefiracetam had no adverse effects on wildtype swimming behavior; however incubation with 125 µM increased the absolute smooth orientation (turning angle as a function of time) of wildtype larvae. Incubation with 85 µM and 125 µM Nefiracetam had no effect on *myo7aa*$^{-/-}$ larvae but incubation with 250 µM did decrease the absolute smooth orientations compared to those untreated. Individual turning angles from populations of wildtype and *myo7aa*$^{-/-}$ larvae treated and untreated were used to construct the lines (t-test). The black bold line represents the mean of the data set and error bars are 95% confidence intervals.

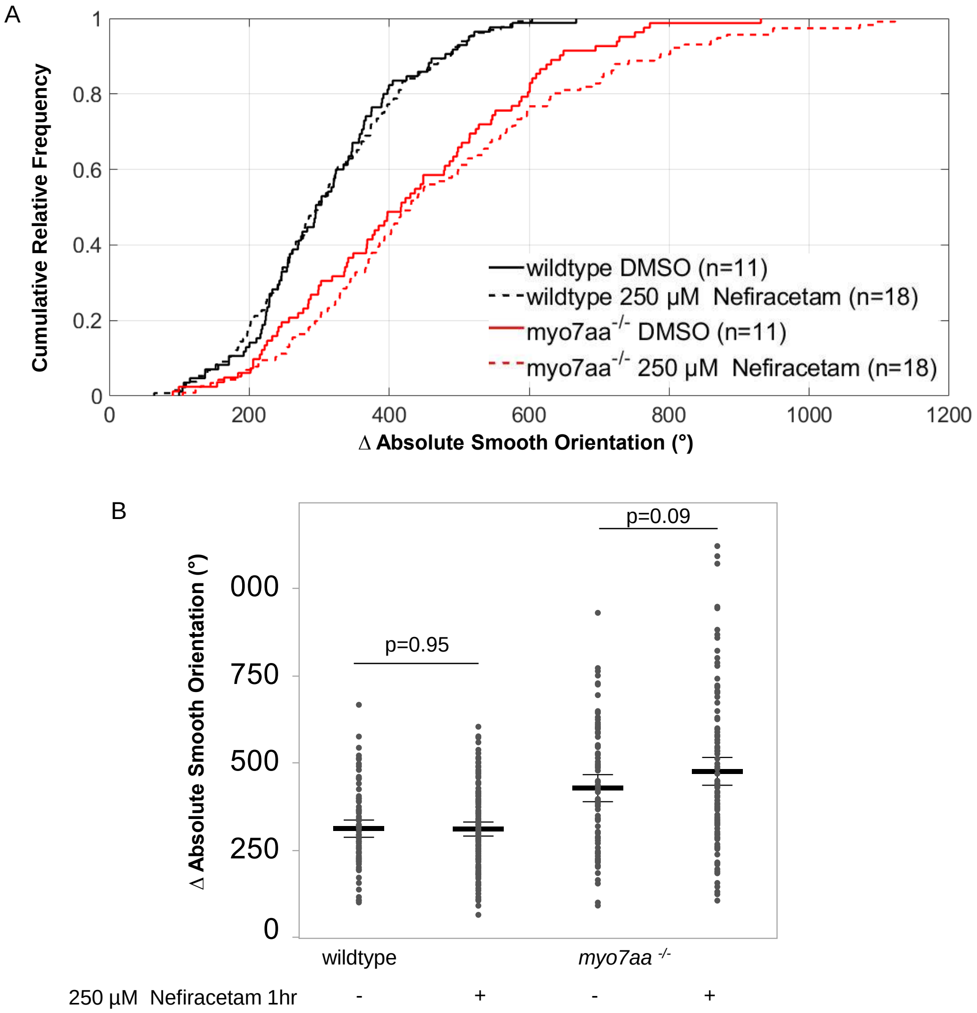


**Fig. S5 1 hour incubation does not suffice for Nefiracetam to affect swimming behavior of wildtype and *myo7aa*$^{-/-}$ mutants.** In order to determine if a shorter incubation period would provide similar responses to an overnight incubation the absolute smooth orientation was quantified. (A, B) Incubation with 250 µM Nefiracetam for 1 hour had no adverse effects on wildtype swimming behavior and did not affect the absolute smooth orientation of the *myo7aa*$^{-/-}$ mutants, indicating that an overnight incubation is required. Individual turning angles from populations of wildtype and *myo7aa*$^{-/-}$ larvae treated and untreated were used to construct the lines (t-test). The black bold line represents the mean of the data set and error bars are 95% confidence intervals.

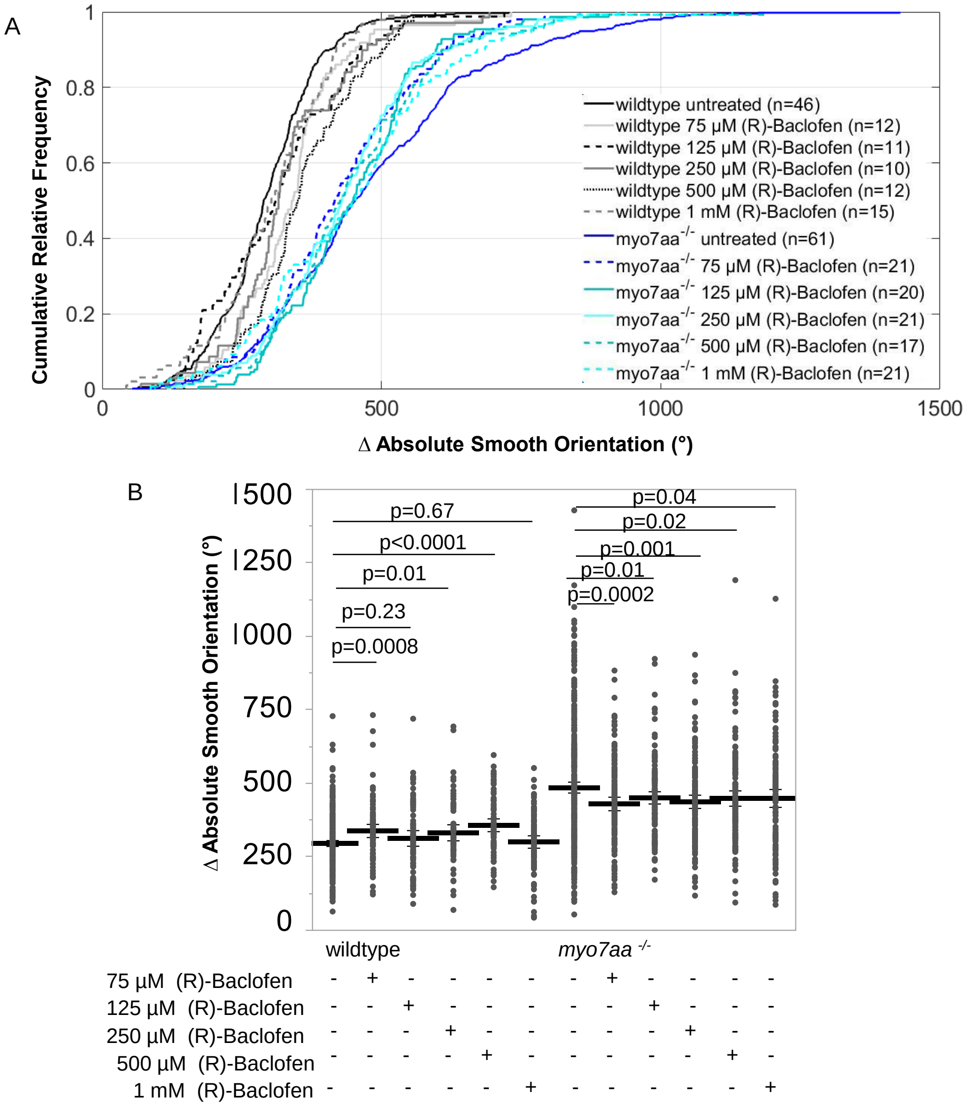


**Fig. S6 Different doses of (R)-Baclofen have different effects on wildtype and *myo7aa*$^{-/-}$ mutant swimming behavior.** (A, B) Incubation with 125 µM and 1 mM (R)-Baclofen had no adverse effects on wildtype swimming behavior; however incubation with 75 µM, 250 µM and 500 µM increased the absolute smooth orientation (turning angle as a function of time) of wildtype larvae. Incubation with all 5 doses of (R)-Baclofen did decrease the absolute smooth orientations of *myo7aa*$^{-/-}$ mutants compared to those untreated. Individual turning angles from populations of wildtype and *myo7aa*$^{-/-}$ larvae treated and untreated were used to construct the lines (t-test). The black bold line represents the mean of the data set and error bars are 95% confidence intervals.

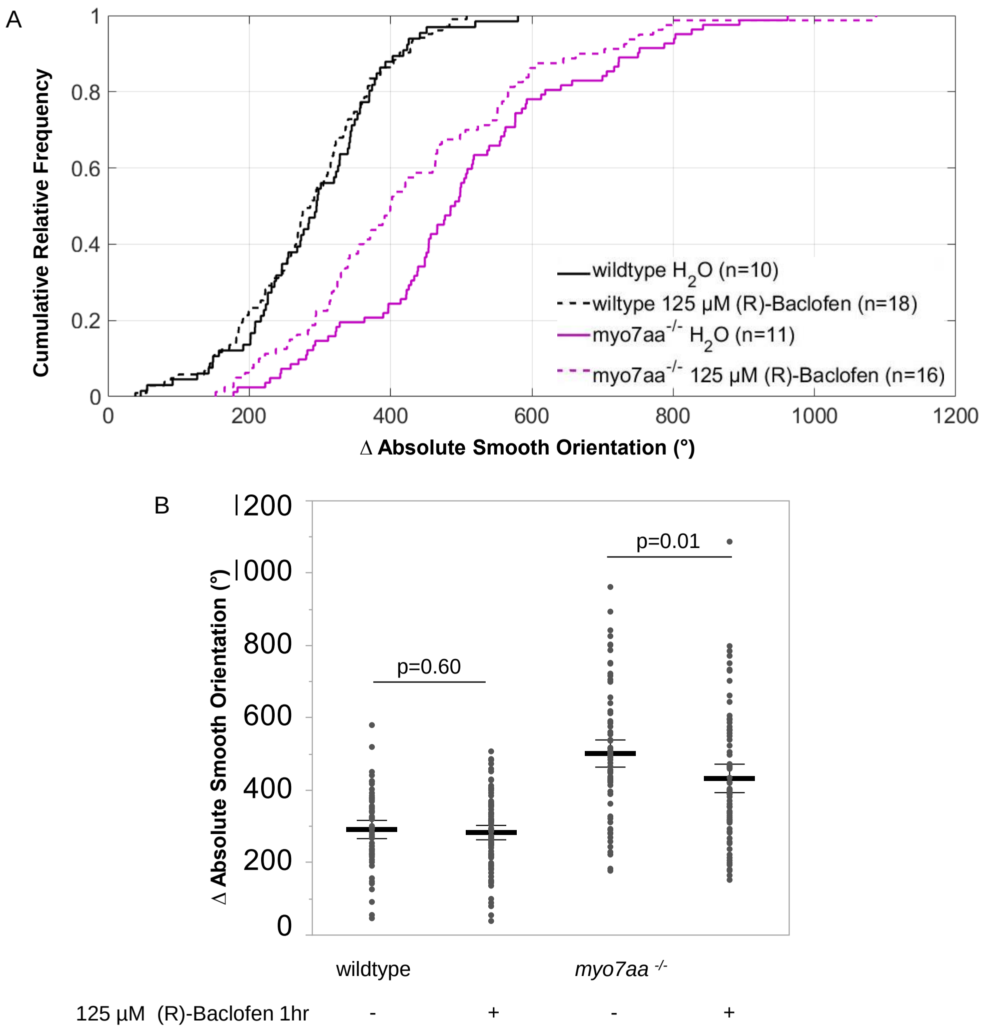


**Fig. S7 1 hour incubation with (R)-Baclofen is sufficient to affect swimming behavior of *myo7aa*$^{-/-}$ mutants.** In order to determine if a shorter incubation period would provide similar responses to an overnight incubation the absolute smooth orientation was quantified. (A, B) Incubation with 125 μM (R)-Baclofen for 1 hour had no adverse effects on wildtype swimming behavior and interestingly did decrease the absolute smooth orientation of the *myo7aa*$^{-/-}$ mutants, indicating that 1 hour incubation is sufficient for this drug. Individual turning angles from populations of wildtype and *myo7aa*$^{-/-}$ larvae treated and untreated were used to construct the lines (t-test). The black bold line represents the mean of the data set and error bars are 95% confidence intervals.

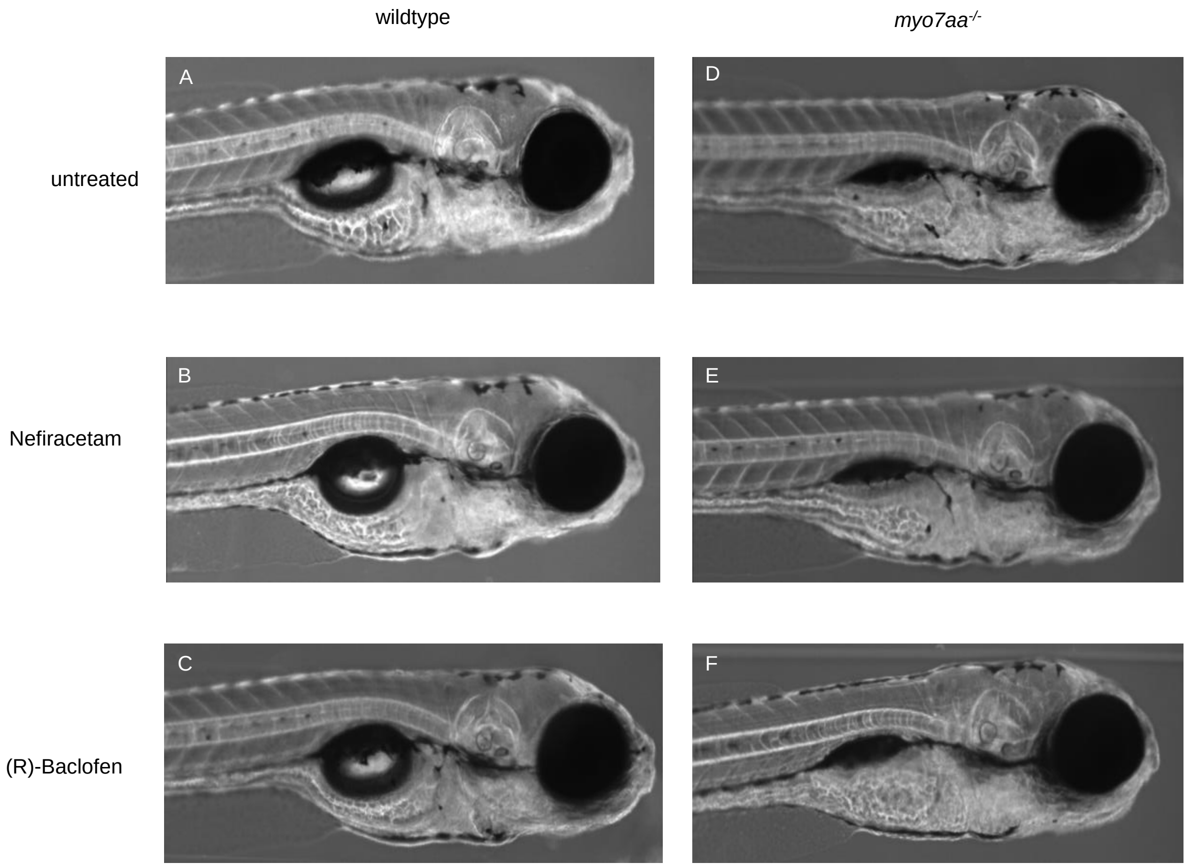


**Fig. S8 250 µM Nefiracetam and 125 µM (R)-Baclofen have no effect on inflation of swim bladder on both wildtype and *myo7aa*$^{-/-}$ larvae.** 4dpf larvae were incubated in either vehicle control, 250 µM Nefiracetam or 125 µM (R)-Baclofen and assessed for any changes in swim bladder. (A, B, C) Wildtype fish swim bladder inflation is unaffected upon incubation with either compound. (D, E, F) *myo7aa*$^{-/-}$ mutant swim bladder inflation is unaffected upon incubation with either 250 µM Nefiracetam or 125 µM (R)-Baclofen.
